# A constitutively monomeric UVR8 photoreceptor allele confers enhanced UV-B photomorphogenesis

**DOI:** 10.1101/2020.08.02.233007

**Authors:** Roman Podolec, Kelvin Lau, Timothée B. Wagnon, Michael Hothorn, Roman Ulm

**Affiliations:** Department of Botany and Plant Biology, Section of Biology, Faculty of Sciences, University of Geneva, 30 Quai E. Ansermet, CH-1211 Geneva 4, Switzerland; Institute of Genetics and Genomics of Geneva (iGE3), University of Geneva, Geneva, Switzerland; Protein Production and Structure Core Facility, École Polytechnique Fédérale de Lausanne, Lausanne, Switzerland

## Abstract

The plant UV-B photoreceptor UVR8 plays an important role in UV-B acclimation and survival. UV-B absorption by homodimeric UVR8 induces its monomerization and interaction with the E3 ubiquitin ligase COP1, leading ultimately to gene expression changes. UVR8 is inactivated through redimerization, facilitated by RUP1 and RUP2. Here, we describe a novel semi-dominant, hyperactive allele, namely *uvr8-17D*, that harbors a glycine-101 to serine mutation. UVR8^G101S^-overexpression led to weak constitutive photomorphogenesis and extreme UV-B responsiveness. UVR8^G101S^ was observed to be predominantly monomeric *in vivo* and, once activated by UV-B, was not efficiently inactivated. Analysis of a UVR8^G101S^ crystal structure revealed the distortion of a loop region normally involved in stabilization of the UVR8 homodimer. Plants expressing a UVR8 variant combining G101S with the previously described W285A mutation exhibited robust constitutive photomorphogenesis. This work provides further insight into UVR8 activation and inactivation mechanisms, and describes a genetic tool for the manipulation of photomorphogenic responses.

## Introduction

Plant growth and development rely on appropriate responses to the light environment. Plants have evolved different photoreceptor families that respond to photons of specific wavelengths: red/far-red light-sensing phytochromes (phyA–E); blue light-sensing cryptochromes (cry1 and cry2), phototropins (phot1 and phot2), and Zeitlupe family proteins (ztl, fkf1, and lkp1); and the UV-B-sensing UV RESISTANCE LOCUS 8 (UVR8) (Rizzini et al., 2011; Tilbrook et al., 2013; Jenkins, 2014; Galvao and Fankhauser, 2015). Photoreceptors shape plant development throughout the entire life cycle. At the seedling stage, photoreceptor-mediated light perception initiates processes such as inhibition of hypocotyl elongation, cotyledon expansion, biogenesis of the photosynthetic machinery, and synthesis of photoprotective pigments (Jenkins, 2017; Yin and Ulm, 2017; Demarsy et al., 2018). Notably, phytochrome, cryptochrome, and UVR8 pathways converge on the CONSTITUTIVELY PHOTOMORPHOGENIC 1–SUPPRESSOR OF PHYA-105 (COP1–SPA) E3 ubiquitin ligase complex, which is responsible for degradation of photomorphogenesis-promoting factors and is inactivated by photoreceptors under light (Hoecker, 2017; Podolec and Ulm, 2018). Perception of UV-B and blue light through UVR8 and cry1, respectively, is crucial for UV-B protection and survival in the field (Rai et al., 2019).

UVR8 is a β-propeller protein that exists in a homodimeric ground state held together by an intricate network of salt-bridge interactions (Rizzini et al., 2011; Christie et al., 2012; Wu et al., 2012; Zeng et al., 2015). Absorption of UV-B photons by specific tryptophan residues that provide chromophore function, of which W285 plays a prime role, disrupts the electrostatic interactions stabilizing the homodimer, resulting in UVR8 monomerization. The active UVR8 monomer interacts with the WD40 domain of COP1 using a co-operative binding mechanism involving the core β-propeller domain of UVR8 together with its disordered C-terminus, which harbors a so-called Valine-Proline (VP) motif (Favory et al., 2009; Cloix et al., 2012; Yin et al., 2015; Lau et al., 2019). Interaction of UVR8 with COP1 inhibits COP1 activity by direct competition with COP1 substrates and remodeling of COP1 E3 ligase complex composition (Huang et al., 2013; Yin and Ulm, 2017; Lau et al., 2019). The COP1 substrate ELONGATED HYPOCOTYL 5 (HY5), a crucial positive regulator of light signaling (Osterlund et al., 2000), is induced transcriptionally and stabilized post-transcriptionally, and is necessary for the regulation of most UVR8-induced genes (Ulm et al., 2004; Brown et al., 2005; Oravecz et al., 2006; Favory et al., 2009; Huang et al., 2012; Binkert et al., 2014). Additionally, UVR8 regulates the stability or DNA-binding activity of a set of transcription factors affecting UV-B responses (Hayes et al., 2014; Liang et al., 2018; Yang et al., 2018; Sharma et al., 2019; Qian et al., 2020; Tavridou et al., 2020a; Tavridou et al., 2020b; Yang et al., 2020).

*REPRESSOR OF UV-B PHOTOMORPHOGENESIS 1* (*RUP1*) and *RUP2* are among early UV-B-induced genes and encode components of a negative feedback loop that downregulates UVR8 signaling (Gruber et al., 2010). RUP1 and RUP2 physically interact with UVR8 and promote UVR8 redimerization back to its inactive state (Gruber et al., 2010; Heijde and Ulm, 2013). Additionally, a role for RUP1 and RUP2 as components of an E3 ligase complex mediating HY5 degradation has recently been suggested (Ren et al., 2019).

To dissect UVR8 function, several studies have made use of targeted mutagenesis to assess the role of individual tryptophan residues in photoreception and the importance of electrostatic interactions in mediating dimerization (Rizzini et al., 2011; Christie et al., 2012; O’Hara and Jenkins, 2012; Wu et al., 2012; Heijde et al., 2013; Huang et al., 2014; Heilmann et al., 2016). For example, UVR8^W285F^ is locked in its inactive homodimeric form and is thus unresponsive to UV-B (Rizzini et al., 2011; Christie et al., 2012; Wu et al., 2012; Heijde et al., 2013). Several other mutant forms, including UVR8^W285A^ and UVR8^D96N,D107N^, display a constitutive interaction with COP1, whereas their downstream effects differ (Heijde et al., 2013; Huang et al., 2014; Heilmann et al., 2016). Notably, UVR8^W285A^ is a weak UVR8 dimer that does not respond to UV-B activation (Rizzini et al., 2011; Christie et al., 2012; Heijde et al., 2013). Overexpression of UVR8^W285A^ leads to constitutive photomorphogenesis, likely through inhibition of COP1 (Heijde et al., 2013). On the other hand, UVR8^D96N,D107N^ is constitutively monomeric; however, despite its constitutive interaction with COP1, UV-B exposure is required to induce physiological responses (Christie et al., 2012; Heilmann et al., 2016).

We present here *uvr8-17D*, a novel semi-dominant, hypersensitive *uvr8* mutant allele that harbors the UVR8^G101S^ protein variant. We found that UVR8^G101S^ is predominantly monomeric *in vivo*, requires UV-B for activation, and confers an exaggerated UV-B response when expressed at wild-type levels, which is attributed to impaired redimerization and sustained UVR8 activity. Moreover, we describe the engineered UVR8^G101S,W285A^ variant with amazingly strong constitutive activity *in vivo*.

## Results

### *uvr8-17D* exhibits enhanced UV-B-induced photomorphogenesis

To identify regulators of the UVR8 pathway, we screened an ethyl methanesulfonate (EMS)– mutagenized Arabidopsis population (Col-0 accession) for mutants with an aberrant hypocotyl phenotype when grown for 4 d under weak white light supplemented with UV-B. Several mutants with long hypocotyls contained mutations in the *UVR8* coding sequence; we thus named these alleles *uvr8-16* and *uvr8-18–30* (Supplementary Fig. 1a). Conversely, several of the mutants displaying short hypocotyls in the screen conditions contained mutations in *RUP2*; we named these novel alleles *rup2-3–10* (Supplementary Fig. 1b). However, one mutant with enhanced UV-B-responsiveness did not carry a *RUP2* mutation and showed a semi-dominant hypocotyl phenotype under UV-B (Supplementary Fig. 2a). Interestingly, in this mutant we found a G to A transition in the *UVR8* locus that results in a glycine-101 to serine (G101S) amino acid change (Fig. 1a); we thus named this novel allele *uvr8-17D*.

**Fig. 1.**
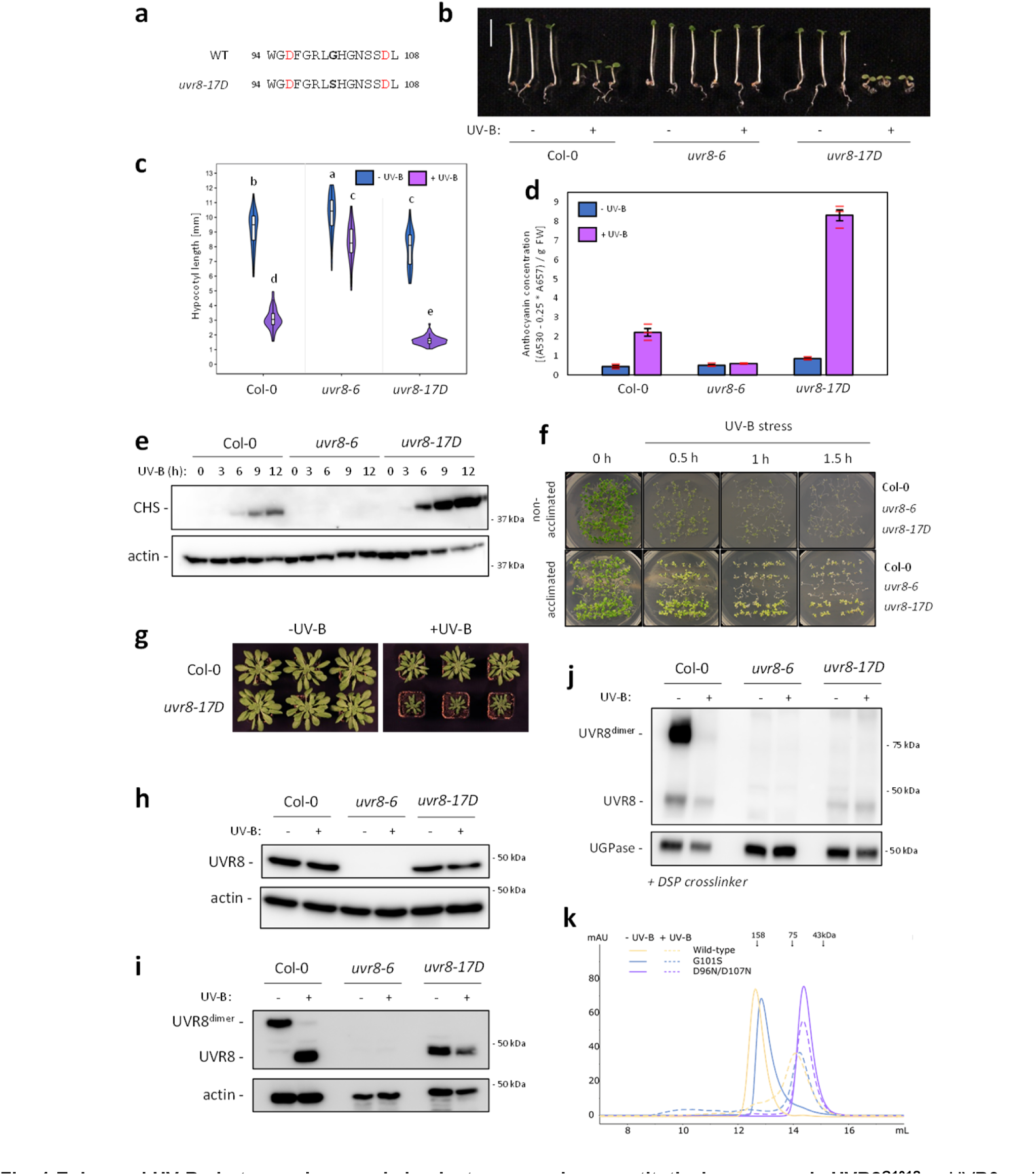
Enhanced UV-B photomorphogenesis in plants expressing constitutively monomeric UVR8^G101S^. **a** UVR8 amino-acid sequence of residues 94–108 in wild type (WT) and *uvr8-17D*. The G101S mutation in *uvr8-17D* is indicated in bold. Asp- 96 and -107 involved in dimer interaction are indicated in red. **b** Representative images of wild-type (Col-0), *uvr8-6* null-mutant, and *uvr8-17D* seedlings grown in white light or white light supplemented with UV-B. Bar = 5 mm. **c** Quantification of hypocotyl length (*N* > 60); shared letters indicate no statistically significant difference in the means (*P* < 0.05). **d** Anthocyanin concentration; values of independent measurements (red bars), means and SEM are shown (*N* = 3). **e** Immunoblot analysis of CHS and actin (loading control) protein levels in Col-0, *uvr8-6*, and *uvr8-17D* grown in white light for 4 d and then white light supplemented with UV-B for 0–12 h. **f** Survival of Col-0, *uvr8-6*, and *uvr8-17D* seedlings after UV-B stress. Seedlings were grown for 7 d in white light (non-acclimated) or white light supplemented with UV-B (acclimated), then exposed to varying durations (0–1.5 h) of broadband UV-B stress. Pictures were taken after a 7-d recovery period. **g** Rosette phenotype of Col-0 and *uvr8-17D* grown for 56 d under short-day conditions in white light or white light supplemented with UV-B. **h** Immunoblot analysis of UVR8 and actin (loading control) protein levels in 7-d-old Col-0, *uvr8-6*, and *uvr8-17D* seedlings. **i** UVR8 dimer/monomer status in Col-0 and *uvr8-17D*. Protein samples were extracted from dark-grown seedlings either exposed or not exposed to 15 min of saturating UV-B and analyzed using immunoblot analysis of samples separated through SDS-PAGE without prior heat denaturation. Actin is shown as loading control. **j** Immunoblot analysis of UVR8 in DSP-crosslinked extracts of dark-grown Col-0, *uvr8-6*, and *uvr8-17D* seedlings either exposed or not exposed to 15 min of saturating UV-B. UGPase is shown as loading control. **k** Size-exclusion chromatography of recombinant UVR8, UVR8^G101S^, and UVR8^D96N,D107N^ proteins purified from Sf9 insect cells, with and without UV-B treatment.

*uvr8-17D* seedlings exhibited shorter hypocotyls and elevated anthocyanin levels under UV-B compared with wild type, whereas under white light there were only minor phenotypic differences (Fig. 1b–d). Hypocotyl lengths were indistinguishable in dark-grown seedlings, suggesting that *uvr8-17D* does not exhibit significant constitutive activity (Supplementary Fig. 2b, c). Consistent with the UV-B-hypersensitive phenotype, CHALCONE SYNTHASE (CHS) and HY5 accumulated to higher levels in *uvr8-17D* than in wild type (Fig. 1e, Supplementary Fig. 2d). *uvr8-17D* seedlings also showed enhanced UV-B acclimation and stress tolerance (Fig. 1f). Compared to wild type, *uvr8-17D* plants showed greater dwarfing at rosette stage under UV-B, whereas they grew normally in the absence of UV-B (Fig. 1g). Altogether, the *uvr8-17D* phenotype and UV-B responsiveness resembled that of UVR8 overexpression (UVR8-OX) lines (Supplementary Fig. 3a–c) (Favory et al., 2009); however, UVR8^G101S^ levels in *uvr8-17D* were comparable to endogenous UVR8 levels in wild type (Fig. 1h, Supplementary Fig. 3c).

We tested whether UVR8^G101S^ is affected in its homodimeric ground state or in UV-B-activated monomerization by SDS-polyacrylamide gel electrophoresis (SDS-PAGE) analysis without sample heat denaturation, as previously described (Rizzini et al., 2011). In this assay, UVR8^G101S^ migrated as a constitutive monomer, whereas wild-type UVR8 derived from seedlings grown in the absence of UV-B or irradiated with saturating UV-B migrated as a homodimer or a monomer, respectively (Fig. 1i). Moreover, in contrast to wild-type UVR8, no UVR8^G101S^ homodimers could be detected *in vivo* by DSP crosslinking (Fig. 1j). However, when assessing the oligomeric state of recombinant UVR8^G101S^ *in vitro*, we found through analytical size-exclusion chromatography experiments that the UVR8^G101S^ mutant protein purified from insect cells behaved similarly to wild-type UVR8, in that it eluted as a dimer under –UV-B conditions and as a monomer after UV-B treatment (Fig. 1k). We thus conclude that UVR8^G101S^ is a hypersensitive UVR8 variant, which is very likely associated with its monomeric state *in vivo*.

### UVR8^G101S^ overexpression leads to extreme UV-B photomorphogenesis

UVR8^G101S^ expression at levels comparable to a previously described UVR8-OX line (Favory et al., 2009) (Fig. 2a) resulted in strikingly enhanced anthocyanin accumulation under UV-B (Fig. 2b, c). Interestingly, hypocotyl elongation was strongly reduced in UVR8^G101S^-OX lines when compared with wild type and UVR8-OX, even in the absence of UV-B (Fig. 2d, e). To determine whether the latter was due to the extreme sensitivity of UVR8^G101S^ to very low levels of UV-B from the fluorescent tubes in this light field (Heijde et al., 2013) or to the intrinsically constitutive activity of overexpressed UVR8^G101S^, we measured the hypocotyl length of seedlings grown in the dark or in monochromatic red or blue light. Although UVR8^G101S^-OX seedlings displayed only a slightly altered hypocotyl-growth phenotype in darkness, some of the lines showed open cotyledons, suggesting that UVR8^G101S^-OX induces weak constitutive photomorphogenesis (Supplementary Fig. 4a, b). More strikingly, comparable to their phenotype described above (Fig. 2d, e), UVR8^G101S^-OX seedlings were indeed shorter in continuous red or blue conditions (Supplementary Fig. 4c–f), suggesting that UVR8^G101S^ broadly sensitizes photomorphogenesis. Consistently, mature UVR8^G101S^-OX plants exhibited dwarfing and early flowering even in the absence of UV-B, with dwarfing further exaggerated under UV-B (Fig. 2f). In agreement with this enhanced photomorphogenesis, UVR8^G101S^-OX lines showed a constitutively enhanced UV-B tolerance that was even further enhanced upon UV-B acclimation (Fig. 2g).

**Fig. 2.**
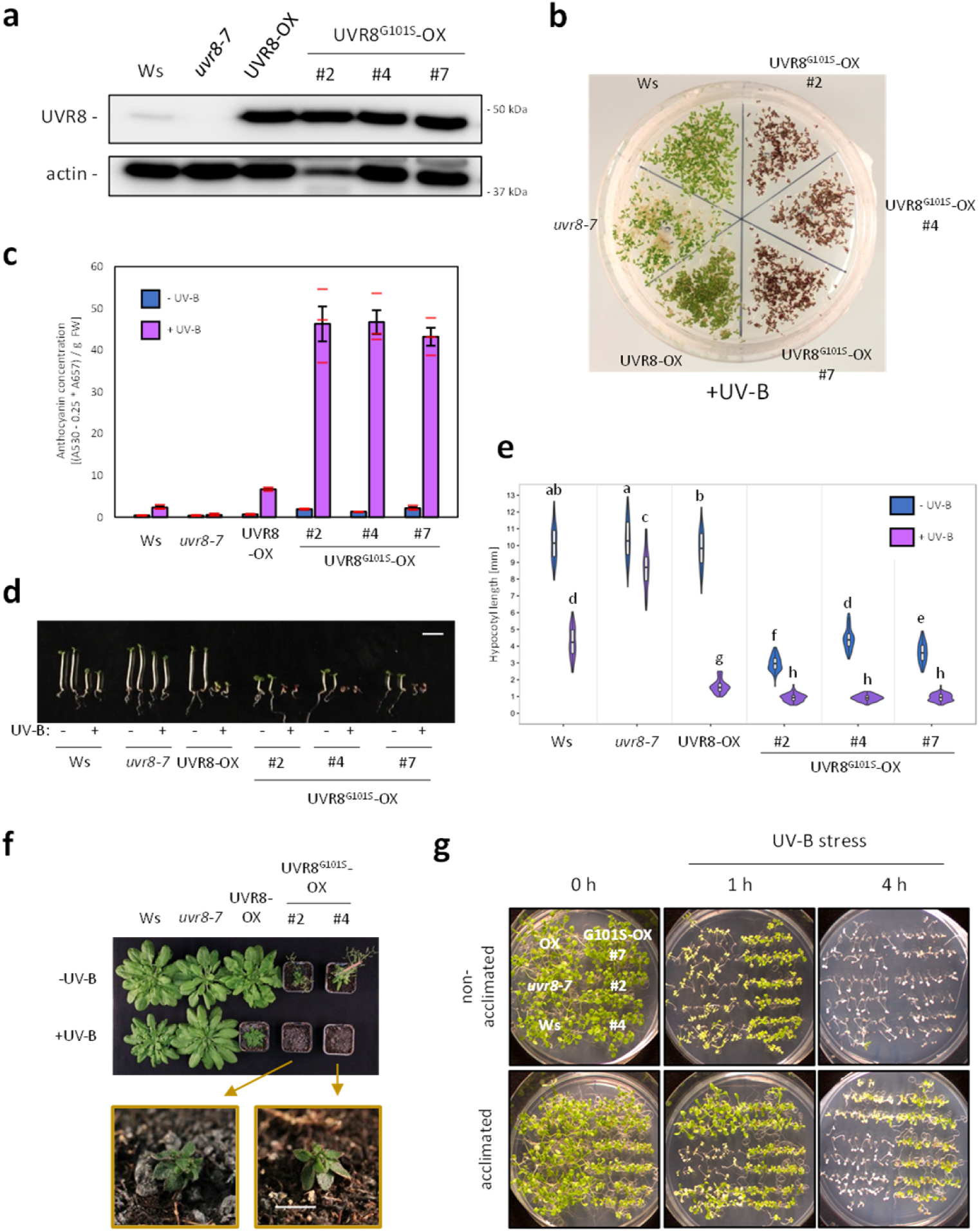
Overexpression of UVR8^G101S^ results in strongly enhanced UV-B photomorphogenesis. **a** Immunoblot analysis of UVR8 and actin (loading control) protein levels in wild type (Ws), *uvr8-7*, *uvr8-7*/Pro_35S_:UVR8 (UVR8-OX), and three independent *uvr8-7*/Pro_35S_:UVR8^G101S^ (UVR8^G101S^-OX #2, #4, and #7) lines. **b** Representative image of plate-grown seedlings grown under UV-B. **c** Anthocyanin concentration of seedlings depicted in **b**. Values of independent measurements (red bars), means, and SEM are shown (*N* = 3). **d** Representative images of seedlings grown in white light or white light supplemented with UV-B. Bar = 5 mm. **e** Quantification of hypocotyl length of seedlings depicted in **d** (*N* > 60). Shared letters indicate no statistically significant difference in the means (*P* < 0.05). **f** Rosette phenotype of representative plants grown for 56 d under short-day conditions in white light or white light supplemented with UV-B. The closeups show UV-B-grown UVR8^G101S^-OX plants. Bar = 5 mm. **g** Survival of seedlings grown for 7 d in white light (non-acclimated) or white light supplemented with UV-B (acclimated) then exposed for varying durations (0–4 h) to UV-B stress. Pictures were taken after a 7-d recovery period.

### UVR8^G101S^ shows no increased binding affinity for COP1 under UV-B

In agreement with the constitutive activity of UVR8^G101S^ when overexpressed, yeast two-hybrid (Y2H) analysis showed that UVR8^G101S^ interacts weakly with COP1 in the absence of UV-B (Supplementary Fig. 5a). UV-B irradiation of the yeast strongly enhanced the UVR8^G101S^–COP1 interaction to a level comparable to the UVR8–COP1 interaction under UV-B, which, however, is strictly UV-B dependent (Supplementary Fig. 5a) (Rizzini et al., 2011). Similarly, in comparison to UVR8, recombinant UVR8^G101S^ showed enhanced *in vitro* binding to the COP1 WD40 domain in the absence of UV-B as shown through quantitative grating-coupled interferometry (GCI) binding assays (Supplementary Fig. 5b). UV-B enhanced the affinity of both UVR8 and UVR8^G101S^ for COP1, but the effect was much stronger for wild-type UVR8 (Supplementary Fig. 5b). Importantly, the G101S mutation did not increase the affinity of UV-B-exposed UVR8^G101S^ for COP1. This suggests that increased affinity of UVR8^G101S^ for COP1 does not underlie the enhanced activity of UVR8^G101S^ *in vivo*. This also agrees with that COP1 activity under UV-B is mainly inhibited through the C-terminal VP motif of UVR8 (Cloix et al., 2012; Yin et al., 2015; Lau et al., 2019), and our observations here that CRISPR/Cas9-generated mutations causing truncation of the UVR8 C-terminus suppressed the UV-B-hypersensitive phenotype of *uvr8-17D* (Supplementary Fig. 6a–d).

### UVR8^G101S,W285A^ shows strong constitutive photomorphogenesis

We hypothesized that combining the G101S mutation conferring UV-B hypersensitivity with the W285A mutation conferring weak constitutive activity (Heijde et al., 2013) would create a strong constitutive UVR8 variant. Thus, we created UVR8^G101S,W285A^, which, in contrast to UVR8^W285A^ and UVR8^G101S^ that were still able to form homodimers *in vitro*, appeared fully monomeric *in vitro* as well as in the crosslinking assay *in vivo* (Fig. 3a, Supplementary Fig. 7a). We observed a strongly elevated constitutive interaction of UVR8^G101S,W285A^ with COP1 compared to the COP1–UVR8^G101S^ or COP1–UVR8^W285A^ interaction in the Y2H system (Fig. 3b) as well as in GCI assays (Fig. 3c, see also Supplementary Fig. 5b). When we attempted to generate transgenic UVR8^G101S,W285A^-OX lines (Pro35S:UVR8^G101S,W285A^), we noticed seedling lethality in the T1 generation, strongly resembling *cop1* null alleles (McNellis et al., 1994). We thus generated lines expressing UVR8^G101S,W285A^ under control of the estradiol-inducible XVE system or the *UVR8* promoter. Indeed, upon estradiol treatment, XVE:UVR8^G101S,W285A^ lines showed an extreme photomorphogenic phenotype in darkness (Fig. 3d,e). In lines with weak expression of UVR8^W285A^ and UVR8^G101S,W285A^ under the native *UVR8* promoter (Fig. 3f), the former (ProUVR8:UVR8^W285A^) showed only a very minor phenotype in the absence of UV-B, similar to in previous reports (O’Hara and Jenkins, 2012; Heijde et al., 2013), whereas the latter (ProUVR8:UVR8^G101S,W285A^) showed a striking constitutive photomorphogenic phenotype resembling that associated with UVR8^W285A^-OX (Fig. 3f–h and Supplementary Fig. 7b,c). Of note, this constitutive photomorphogenesis was in spite of UVR8^G101S,W285A^ protein levels being lower than endogenous UVR8 in wild type (Fig. 3f). We conclude that the combination of G101S and W285A in UVR8 yields a fully monomeric variant with very strong constitutive activity *in vivo*.

**Fig. 3.**
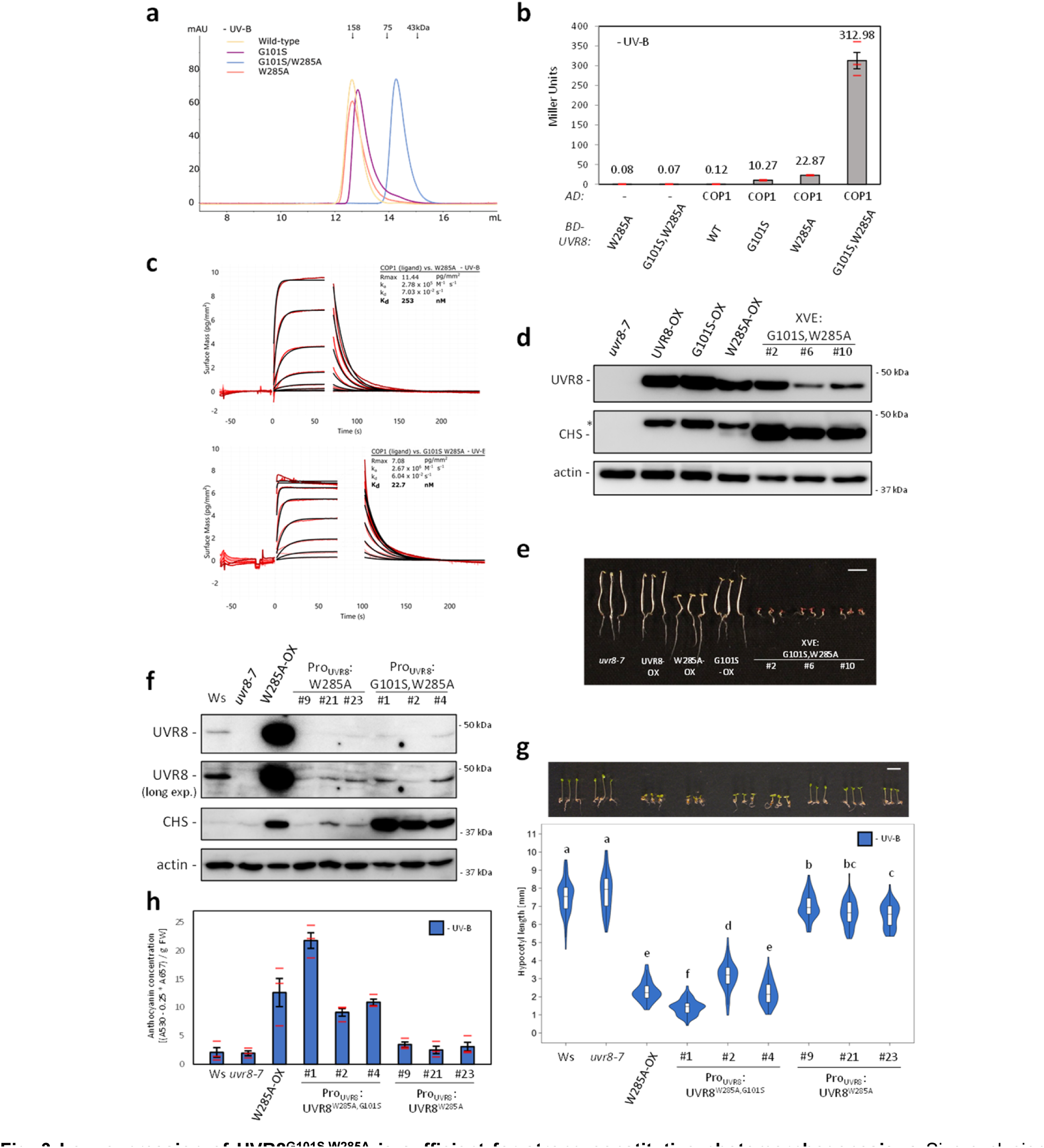
Low expression of UVR8^G101S,W285A^ is sufficient for strong constitutive photomorphogenesis. **a** Size-exclusion chromatography of recombinant UVR8 (Wild-type), UVR8^G101S^ (G101S), UVR8^W285A^ (W285A), and UVR8^G101S,W285A^ (G101S/W285A) proteins purified from Sf9 insect cells. **b** Quantitative Y2H analysis of the interaction between COP1 and UVR8 (WT), UVR8^G101S^, UVR8^W285A^, and UVR8^G101S,W285A^ in the absence of UV-B. AD, activation domain; BD, DNA binding domain. **c** Binding kinetics of the full-length UVR8^W285A^ and UVR8^G101S,W285A^ versus the COP1 WD40 domain obtained by GCI experiments. Sensorgrams of protein injected are shown in red, with their respective heterogenous ligand binding model fits in black. The following amounts were typically used: ligand – COP1 (2,000 pg/mm^2^); analyte – UVR8 (2 μM highest concentration). ka = association rate constant, kd = dissociation rate constant, Kd = dissociation constant. **d** Immunoblot analysis of UVR8, CHS and actin (loading control) protein levels in seedlings of *uvr8-7*, *uvr8-7*/Pro_35S_:UVR8 (UVR8-OX), *uvr8-7*/Pro_35S_:UVR8^G101S^ #2 (G101S-OX), *uvr8-7*/Pro_35S_:UVR8^W285A^ (W285A-OX), and three independent lines of *uvr8-7*/XVE:UVR8^G101S,W285A^ (XVE:G101S,W285A #2, #6, and #10) grown on 5 μM estradiol. Asterisk = residual UVR8 signal after stripping of the PVDF membrane. **e** Representative image of seedlings described in **d** grown for 4 d in darkness on plates supplemented with 5 μM estradiol. Bar = 5 mm. **f** Immunoblot analysis of UVR8, CHS, and actin (loading control) protein levels in wild type (Ws), *uvr8-7*, *uvr8-7*/Pro_35S_:UVR8^W285A^, and three independent lines of each of *uvr8-7*/Pro_UVR8_:UVR8^W285A^ (#9, #21, and #23) and *uvr8-7*/Pro_UVR8_:UVR8^G101S,W285A^ (#1, #2, and #4). **g** Representative seedling images and quantification of hypocotyl length (*N* > 60) for seedlings described in **f**. Shared letters indicate no statistically significant difference in the means (*P* < 0.05). Bar = 5 mm. **h** Anthocyanin concentration of seedlings described in **f**. Values of independent measurements (red bars), means, and SEM are shown (*N* = 3).

### The combination of G101S and W285F confirms a basal constitutive activity for UVR8^G101S^

We further generated and characterized a UVR8^G101S,W285F^ variant. The W285F mutation was expected to abolish all UV-B responsiveness (Rizzini et al., 2011; Heijde et al., 2013), whereas, as described above for UVR8^G101S^-OX (Fig. 2c–e), the G101S mutation results in weak constitutive activity in conditions devoid of UV-B. Integrating the G101S mutation into the constitutively dimeric UVR8^W285F^ variant caused strongly reduced dimer stability in UVR8^G101S,W285F^ (Supplementary Fig. 7a). As predicted, UVR8^G101S,W285F^ retained a basal interaction with COP1 in the Y2H system, similar to COP1–UVR8^G101S^ interaction in the absence of UV-B; however, by contrast, COP1–UVR8^G101S,W285F^ interaction was not further enhanced under UV-B (Supplementary Fig. 8a). UVR8^G101S,W285F^-OX lines exhibited a phenotype in both the absence and presence of UV-B resembling UVR8^G101S^-OX lines in –UV-B conditions (Supplementary Fig. 8b–e), confirming that the phenotype induced by UVR8^G101S^ in the absence of UV-B is not linked to W285-mediated photoactivation.

### The G101S mutation distorts a critical loop interaction in the UVR8 dimer

We next investigated the effects of the G101S mutation in structural detail. No crystals could be obtained for the UVR8^G101S^ protein, yet crystals developed for UVR8^G101S,W285A^, diffracting to 1.75 Å resolution (Supplementary Table 1). The structure revealed a non-symmetric dimeric arrangement of UVR8^G101S,W285A^ in the crystal lattice, as the relative position of the monomers are rotated ∼10° with respect to the previously determined symmetric UVR8 wild-type dimer (Supplementary Fig. 9, Supplementary Fig. 10) (Christie et al., 2012). The altered dimeric assembly of UVR8^G101S,W285A^ contains a unique salt-bridge/hydrogen bond network between the two monomers with a reduced number of interactions compared to the wild-type dimer (Supplementary Fig. 10, Supplementary Table 2). The G101S mutation maps to a loop that forms part of the crystallographic UVR8 dimer interface (Fig. 4a, Supplementary Fig. 11). A structural superposition with wild-type UVR8 revealed that the position of the loop in UVR8^G101S,W285A^ is distorted by the G101S mutation (Fig. 4b–d). Importantly, the slightly larger Ser side-chain cannot be accommodated within the loop. This forces a movement of the loop that affects the position of Asp-96 and -107, the mutation of which to Asn has been previously shown to impair UVR8 homodimerisation (Christie et al., 2012; Heilmann et al., 2016). The W285A mutation results in rearrangements of adjacent Trp side-chains reminiscent to the movements observed in the UVR8^W285A^ crystal structure (Wu et al., 2012), highlighting the potential changes upon UV-B irradiation (Fig. 4c). We also obtained crystal structures of UVR8^D96N,D107N^ and UVR8^D96N,D107N,W285A^ variants that revealed no large structural rearrangements in the loop (Fig. 4e,f, Supplementary Fig. 12). Together, our structural analysis reveals that the G101S mutation affects the conformation of a loop region involved in UVR8 homodimer stabilization.

**Fig. 4.**
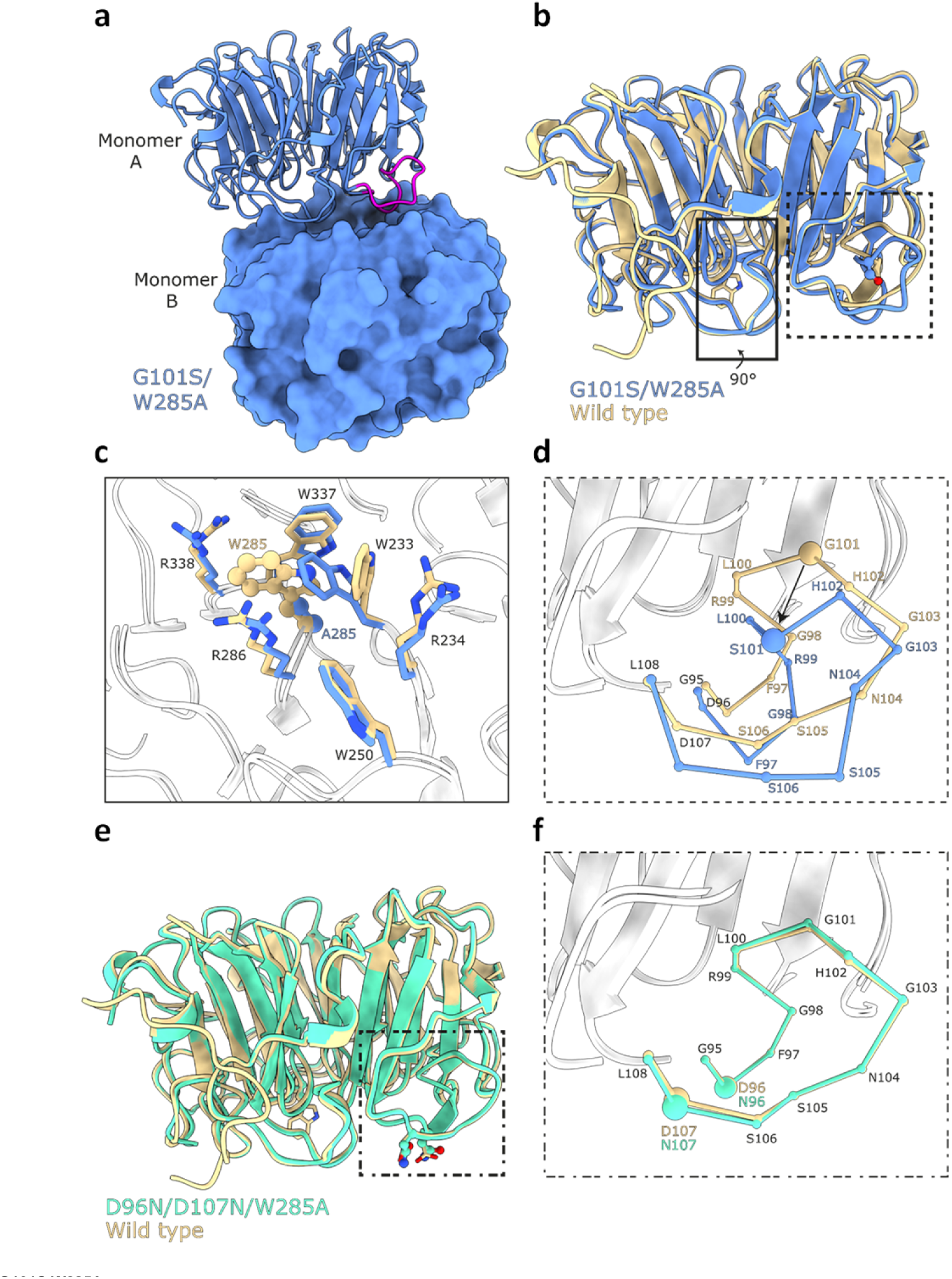
UVR8^G101S,W285A^ distorts a critical interaction loop at the dimer interface. **a** Ribbon and surface representations of the UVR8^G101S,W285A^ dimer. Highlighted in magenta is the critical loop containing the G101S mutation. **b** Superposition of UVR8^G101S,W285A^ (blue) with a wild-type UVR8 (PDB:4D9S, yellow) in ribbon representation. The site of mutation, residue 101, is represented in a ball-and-stick manner and colored by their atomic identity. Residue 285 is represented by sticks. **c** Zoomed-in view of the site of the W285A mutation. The site of mutation is represented in a ball-and-stick manner and the surrounding residues are shown as sticks. **d** Zoomed-in view of the loop containing the G101S mutation. The loop is represented with each ball corresponding to a C_α_ carbon to highlight the structural rearrangements of the loop. **e** Superposition of UVR8^D96N,D107N,W285A^ (green) with UVR8 represented as in **b**. **f** Zoomed-in view of the loop containing G101 represented as in **d**.

### Enhanced activity of UVR8^G101S^ is caused by impaired inhibition through RUP1 and RUP2

Mutations in *RUP1* and *RUP2* enhance the UV-B photomorphogenic phenotype, as active monomeric UVR8 is more prevalent in *rup1 rup2* (Gruber et al., 2010; Heijde and Ulm, 2013). We hypothesized that monomeric UVR8^G101S^ enhances UV-B signaling due to its impaired redimerization, thus mimicking a *rup1 rup2* mutant. We first tested the interaction of UVR8 and UVR8^G101S^ with RUP2 in the Y2H system. Overall, the UVR8–RUP2 interaction was only slightly affected in yeast strains containing UVR8^G101S^ compared to UVR8 (Fig. 5a).

**Fig. 5.**
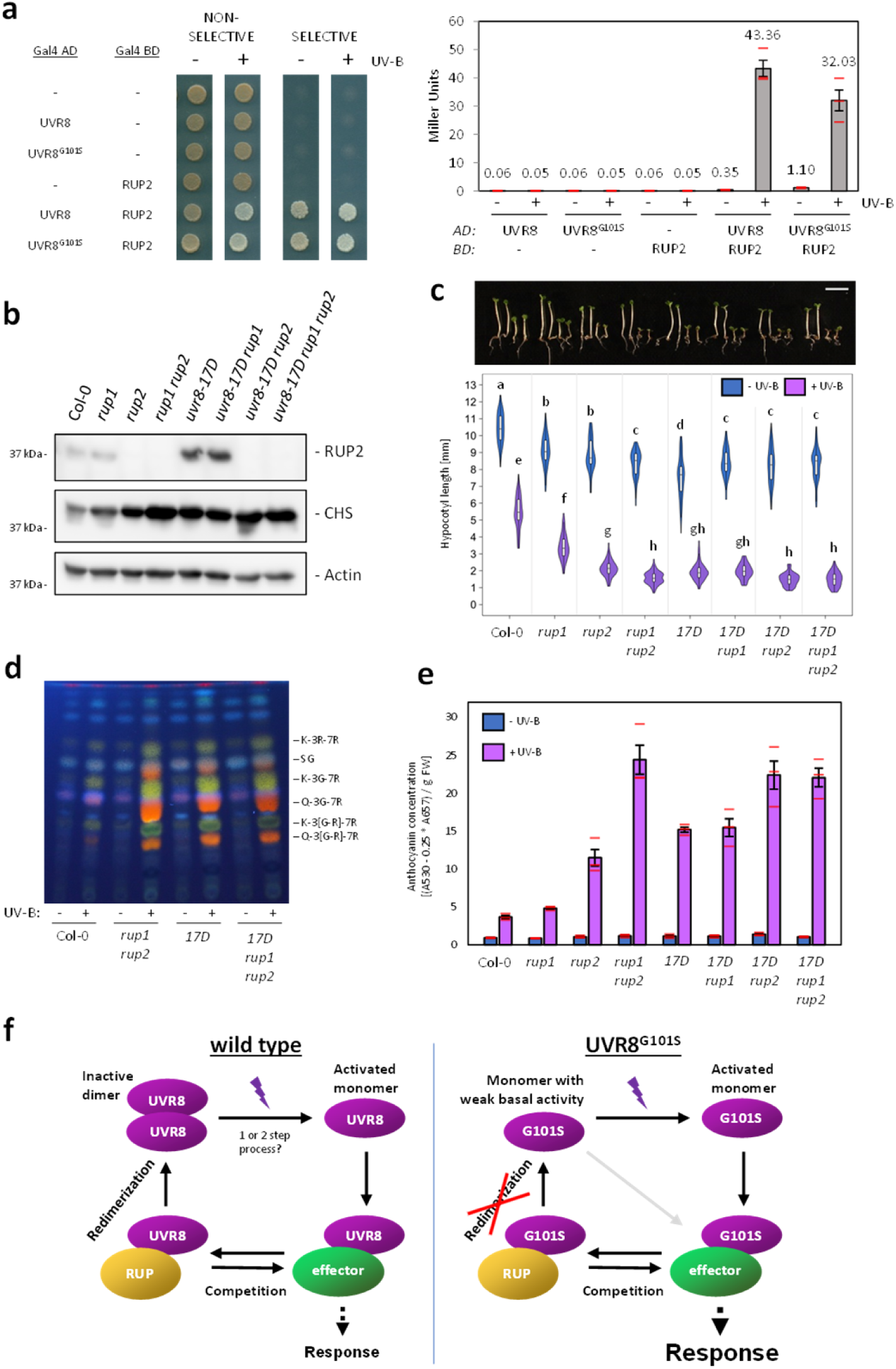
The enhanced activity of UVR8^G101S^ depends on the presence of RUP1 and RUP2. **a** Y2H analyses of the interactions of RUP2 with UVR8 and UVR8^G101S^ in the presence or absence of UV-B. Left: growth assay on selective SD/-Trp/-Leu/-His medium. Right: quantitative β-galactosidase assay. AD, activation domain; BD, DNA binding domain. **b** Immunoblot analysis of RUP2, CHS, and actin (loading control) protein levels in seedlings of wild type (Col-0), *rup1*, *rup2*, *rup1 rup2*, *uvr8-17D*, *uvr8-17D rup1*, *uvr8-17D rup2*, and *uvr8-17D rup1 rup2* grown for 4 d in weak white light supplemented with UV-B. **c** Representative images of seedlings described in **b** (*17D* = *uvr8-17D*) and quantification of hypocotyl length (*N* > 60). Shared letters indicate no statistically significant difference in the means (*P* < 0.05). Bar = 5 mm. **d** HPTLC analysis of the flavonol glycoside levels in 4-d-old seedlings of wild type (Col-0), *rup1 rup2*, *uvr8-17D*, and *uvr8-17D rup1 rup2* grown in white light or white light supplemented with UV-B. Identified compounds include: K-3R-7R, kaempferol-3-O-rhamnoside-7-O-rhamnoside; SG, sinapoyl glucose; K-3G-7R, kaempferol-3-O-glucoside-7-O-rhamnoside; Q-3G-7R, quercetin-3-O-glucoside-7-O-rhamnoside; K-3[G-R]-7R, kaempferol 3-O-[rhamnosyl-glucoside]-7-O-rhamnoside; Q-3[G-R]-7R, quercetin 3-O-[rhamnosyl-glucoside]-7-O-rhamnoside. **e** Anthocyanin concentration of seedlings described in **b**. Values of independent measurements (red bars), means, and SEM are shown (*N* = 3). **f** Working model of the UVR8 photocycle in the case of wild-type (left) and G101S-mutated UVR8 (right). In the wild type, UV-B induces monomerization of dimeric UVR8 in a one- or two-step photon absorption process. The activated monomer then interacts with effector proteins such as COP1 and transcription factors to induce a photomorphogenic response. RUP1/RUP2 (RUP) proteins compete with these effectors to abrogate signaling activity. Afterwards, RUP-bound UVR8 undergoes redimerization and is brought back to its initial inactive state. UVR8^G101S^ exists as a monomer *in vivo* and exhibits weak constitutive activity in the absence of UV-B, as seen in overexpression lines. UV-B absorption fully activates the UVR8^G101S^ monomer and RUP proteins still compete with other UVR8 signaling effectors. However, the full cycle cannot be completed because redimerization is intrinsically impossible. This results in sustained UVR8 signaling, leading to enhanced UV-B photomorphogenesis.

We then constructed higher-order mutant combinations between *uvr8-17D*, *rup1*, and *rup2*. Under UV-B, RUP2 was detected at an elevated level in *uvr8-17D* and *uvr8-17D rup1* when compared to that in wild type and *rup1*, respectively (Fig. 5b). Interestingly, *uvr8-17D* was similar to *rup1 rup2* in terms of hypocotyl length and flavonol accumulation (Fig. 5c,d), whereas *uvr8-17D* seedlings displayed an intermediate phenotype between a *rup2* and a *rup1 rup2* mutant regarding anthocyanin accumulation (Fig. 5e). Together with the observation that the anthocyanin phenotypes of *uvr8-17D rup2* and *uvr8-17D rup1 rup2* are stronger than that of the single *uvr8-17D* mutant (Fig. 5e), this suggests that RUP2 is still able to partially inhibit _UVR8_G101S.

Interestingly, in the presence of functional RUP1 and RUP2, the UVR8^G101S^ mutation significantly enhances the UV-B phenotype (*uvr8-17D* versus wild type; Fig. 5c,e); however, in their absence, UVR8^G101S^ does not induce stronger UV-B signaling compared to UVR8 (*uvr8-17D rup1 rup2* versus *rup1 rup2*; Fig. 5c,e). Together, this demonstrates that whereas RUP1 and RUP2 proteins retain some ability to negatively regulate UVR8^G101S^, the enhanced activity of this variant is only evident in a background where RUP1 and RUP2 are present. We conclude that the enhanced activity of UVR8^G101S^ is due to impaired redimerization by RUP1 and RUP2, as UVR8^G101S^ shows a strongly reduced ability to dimerize *in vivo*.

## Discussion

*uvr8-17D* is a UV-B-hypersensitive UVR8 photoreceptor allele that is linked with a single glycine-101 to serine amino acid change. The UV-B hypersensitivity of *uvr8-17D* is associated with its monomeric conformation *in vivo*, suggesting that re-dimerization facilitated by RUP1 and RUP2 is impaired and that UVR8^G101S^ therefore remains active much longer than wild-type UVR8 (Fig. 5f). In agreement, *uvr8-17D* phenotypes under UV-B resemble those of *rup1 rup2* double mutants, further supporting the prime importance of UVR8 redimerization to optimally balance UV-B-induced photomorphogenesis with plant growth and development (Gruber et al., 2010; Heijde and Ulm, 2013; Findlay and Jenkins, 2016; Liao et al., 2020; Tissot and Ulm, 2020).

Intriguingly, UVR8^G101S^ appeared monomeric *in vivo*, whereas *in vitro* it appeared monomeric in a gel-based assay and dimeric in a size-exclusion chromatography experiment. The SDS-PAGE assay is not sensitive enough to recognize weak dimeric variants such as UVR8^W285A^, which appears monomeric (Rizzini et al., 2011; Christie et al., 2012; Wu et al., 2012; Heijde et al., 2013). However in crosslinking and size-exclusion chromatography assays, UVR8^W285A^ is indeed dimeric, whereas UVR8^G101S^ crosslinked as a dimer only poorly and only when overexpressed. UVR8^G101S^ was however dimeric in size-exclusion chromatography assays, which could be linked to the different concentrations of UVR8 protein used and the different chemical environments between the *in vitro* assay and the *in vivo* cellular environment.

The monomeric nature of UVR8^G101S^ *in vivo* is reminiscent of the UVR8^D96N,D107N^ variant previously generated based on structural information (Supplementary Fig. 13a) (Christie et al., 2012; Heilmann et al., 2016). Similar to UVR8^G101S^, UVR8^D96N,D107N^ also shows weak constitutive COP1 binding (Supplementary Fig. 13b–e) and requires UV-B for functional activation (Heilmann et al., 2016). By contrast though, no UV-B hypersensitivity was previously reported for transgenic lines expressing UVR8^D96N,D107N^ (Heilmann et al., 2016). However, in our phenotypic assays, transgenic UVR8^D96N,D107N^-OX lines phenocopied the UV-B-hypersensitive phenotype of UVR8^G101S^-OX (Supplementary Fig. 13f,g), suggesting that UVR8^D96N,D107N^ and UVR8^G101S^ indeed have similar enhancing effects on UV-B responsiveness *in vivo*, consistent with our conclusion that the *uvr8-17D* phenotype is directly linked to its monomeric conformation *in vivo*. In support, our atomic structure of UVR8^G101S,W285A^ suggests a direct mechanistic link between the mutation of G101 and the misorientation of the D96 and D107 residues.

The effects of UVR8^G101S^ and UVR8^D96N,D107N^ thus suggest that monomeric UVR8 variants require UV-B to trigger the UV-B signaling pathway, confirming that engineered constitutive UVR8 monomers are not sufficient for strong constitutive UVR8 activity. This confirms that monomerization and activation of UVR8 by UV-B can clearly be uncoupled, at least for mutated UVR8 variants (Heilmann et al., 2016). However, in these cases, the mutant variants are simply monomeric versions that have not been UV-B activated. Whether wild-type UVR8 proteins that monomerize in response to UV-B—a process associated with broader structural changes that likely also affect the VP-containing C-terminus (Heilmann et al., 2015; Yin et al., 2015; Zeng et al., 2015; Camacho et al., 2019; Lau et al., 2019)—can or even need to be similarly UV-B activated remains to be shown. Nonetheless, it may be that the activation of wild-type UVR8 is a two-step process, with photon absorption events required both for chromophore excitation that leads to monomerization and for functional activation. An engineered monomeric UVR8 mutant may be primed for activation by bypassing the need for monomerization prior to activation. Alternatively, one photon absorption event may be sufficient for both monomerization and activation. In this case, monomerization and activation by UV-B would be intrinsically linked in wild-type UVR8. Another possibility is that monomeric wild-type UVR8 absorbs additional UV-B photons as an efficient way to keep the photoreceptor active, without going through redimerization and reactivation by monomerization. Conclusive evidence to address these possibilities will require structural determination of full-length UV-B-activated monomer versus an engineered monomeric UVR8.

Despite remaining monomeric *in vivo*, UVR8^G101S^ retains an interaction with and remains weakly negatively regulated by RUP1 and RUP2. The UVR8–COP1 interaction may be inhibited by direct interaction of RUP proteins with the VP-containing C-terminus of UVR8, which is the same domain that UVR8 uses to bind both COP1 and direct transcription factor targets (Cloix et al., 2012; Yin et al., 2015; Liang et al., 2018; Yang et al., 2018; Lau et al., 2019; Liang et al., 2019). This implies that UVR8 redimerization may proceed via two steps: (i) RUPs outcompete COP1 and other VP-domain interactors, separating them from UVR8, and (ii) RUPs facilitate UVR8 redimerization. As redimerization occurs at a normal rate in a *cop1* background (Heijde and Ulm, 2013), competitive removal of COP1 alone is evidently not sufficient to induce redimerization. In *uvr8-17D*, step (i) could explain the remaining partial activity of RUPs on UVR8^G101S^, whereas step (ii) is abolished.

Importantly, the *uvr8-17D rup1 rup2* triple mutant did not show phenotypic differences compared with *rup1 rup2*, suggesting the mechanisms underlying UVR8^G101S^ hyperactivity are linked to RUP1/RUP2 activity. As a result of more prevalent active UVR8 monomer, the increased signaling induced by UVR8^G101S^ originates early and upstream in the pathway and all subsequent UVR8 signaling mechanisms, including nuclear accumulation, inhibition of COP1 E3 ligase activity, and inhibition of transcription factor activity, are likely to be indirectly promoted by the relatively increased activity of UVR8^G101S^.

UVR8 is widely conserved across the green lineage, and several studies have confirmed its role in mediating UV-B responses in a diverse range of species (Allorent et al., 2016; Tilbrook et al., 2016; Clayton et al., 2018; Li et al., 2018; Soriano et al., 2018; Han et al., 2019; Kondou et al., 2019; Tokutsu et al., 2019; Liu et al., 2020). G101 is universally conserved in UVR8 orthologs, as are residues involved in photoreception and dimer stability (Rizzini et al., 2011; Fernandez et al., 2016; Han et al., 2019). The phenotype of *uvr8-17D* holds the exciting possibility that a homologous glycine-to-serine mutation leads to enhanced UV-B responses in other species as well. This may eventuate as a convenient way to generate UV-B-hypersensitive phenotypes, including enhanced acclimation and UV-B tolerance, made possible by a single nucleotide substitution. Additionally, expressing UVR8^G101S,W285A^ at very low levels is sufficient to produce a striking constitutive photomorphogenic phenotype. Notwithstanding the use of UVR8^G101S^ in other plant species, both for fundamental research and potential biotechnological applications, *uvr8-17D* may serve as a genetic tool to further investigate UVR8 activity regulation and the effects of enhanced UVR8 signaling in Arabidopsis.

## Methods

### Plant material

*Arabidopsis thaliana* mutants *uvr8-6* (Favory et al., 2009), *cop1-4* (Deng et al., 1992), *hy5-215* (Oyama et al., 1997), *rup1-1*, *rup2-1*, and *rup1-1 rup2-1* (Gruber et al., 2010) are all in the Columbia (Col) accession. *uvr8-7*, *uvr8-7*/Pro35S:UVR8 line #2 (UVR8-OX) (Favory et al., 2009), *uvr8-7*/Pro35S:UVR8^W285A^ line #3, and *uvr8-7*/Pro35S:UVR8^W285F^ line #1 (Heijde et al., 2013) are in the Wassilewskija (Ws) accession. Novel *uvr8* and *rup2* alleles were identified within an EMS-mutagenized population in the Col accession.

Combinatorial mutants *uvr8-17D rup1-1*, *uvr8-17D rup2-1*, and *uvr8-17D rup1-1 rup2-1* were generated by genetic crossing. Genotyping of *rup1-1* and *rup2-1* alleles was performed as described (Gruber et al., 2010), and *uvr8-17D* genotyping was through PCR amplification of a genomic fragment with a forward (5ʹ-TCG GGA TGA GAT GAT GAC-3ʹ) and a reverse primer (5ʹ-TAG ACC CAA CAT TGA CCC-3ʹ) followed by digestion with *Hae*III (NEB) yielding diagnostic fragments of 344/173/145 bp for wild type and 489/173 bp for *uvr8-17D*.

To generate transgenic lines expressing UVR8 variants, mutations corresponding to G101S, W285A, W285F, and D96N/D107N were introduced into the *UVR8* coding sequence in pDONR207 (Thermo Fisher Scientific) by site-directed mutagenesis. *UVR8^G101S^*, *UVR8^G101S,W285A^*, *UVR8^G101S,W285F^*, and *UVR8^D96N,D107N^* sequences were inserted into the Gateway-compatible binary vector pB2GW7 (Karimi et al., 2002). *UVR8^G101S,W285A^* was also inserted into the Gateway-compatible binary vector pMDC7 allowing estradiol-inducible expression (Curtis and Grossniklaus, 2003).

To generate transgenic lines expressing UVR8 under its own promoter, the *UVR8* promoter (1,809 bp upstream of the translational start ATG) was respectively cloned upstream of the *UVR8^W285A^* and *UVR8^G101S,W285A^* coding sequences and introduced into the Gateway-compatible binary vector pGWB501 (Nakagawa et al., 2007).

The CRISPR/Cas9 system was used to delete the UVR8 C-terminus in wild type and *uvr8-17D*. An sgRNA directed against the *UVR8* sequence was inserted into the pHEE401E vector (Wang et al., 2015) using overlapping complementary oligos 5ʹ-ATT G*GC GAC ACC CAG CTT TTC CC*-3ʹ and 5ʹ-AAA C*GG GAA AAG CTG GGT GTC GC*-3ʹ, and mutants were identified in T1 after sequencing the corresponding part of the *UVR8* locus.

Arabidopsis plants were transformed using the floral dip method (Clough and Bent, 1998) and transgenic lines with a single insertion locus (75% resistance in T2) were selected for homozygosity in T3.

### Growth conditions

For hypocotyl length measurements, quantification of anthocyanins, analysis of flavonol glycosides, and acclimation assays, seedlings were grown under aseptic conditions on half-strength MS medium (Duchefa) supplemented with 1% [w/v] sucrose (Applichem). For immunoblot analysis, seedlings were grown on sterile half-strength MS medium (Duchefa) without sucrose. For induction of estradiol-inducible expression under the XVE system, the growth medium was supplemented with 5 μM of β-estradiol (Sigma).

After sowing, seeds were stratified for 2 d at 4°C in the dark before the start of the light treatments, which were performed at 22°C as described before (Oravecz et al., 2006; Favory et al., 2009). White light (3.6 μmol m^-2^ s^-1^) was supplied by Osram L18W/30 tubes, which were supplemented with narrowband Philips TL20W/01RS tubes (1.5 μmol m^-2^ s^-1^) for UV-B treatment. To analyze etiolated growth in darkness, seeds were exposed to 6-h white light (approx. 60 μmol m^-2^ s^-1^) to induce germination before being transferred back to darkness. For monochromatic light treatments, a light-emitting diode (LED) chamber (floraLEDs, CLF Plant Climatics) was used with 5 μmol m^-2^ s^-1^ of blue light or 30 μmol m^-2^ s^-1^ of red light.

For analysis of vegetative growth, plants were grown in short days (8-h/16-h light/dark cycles) in UV-B lamp–containing GroBanks (CLF Plant Climatics), under white light (120 μmol m^-2^ s^-1^) or white light supplemented with UV-B (1.5 μmol m^-2^ s^-1^) (Arongaus et al., 2018).

For broadband UV-B treatments (acclimation assays, analyses of dimer/monomer status), Philips TL40W/12RS UV-B tubes were used (Favory et al., 2009; Rizzini et al., 2011).

### Hypocotyl length measurements

Quantification of hypocotyl length was performed as described previously (Lau et al., 2019). At least 60 seedlings for each genotype and experimental condition were aligned on plates, which were then scanned and hypocotyl length was measured using the NeuronJ plugin of ImageJ (Meijering et al., 2004). Violin and box plots were generated using the ggplot2 package in R (Wickham, 2009), and statistically different groups were determined using the Tukey HSD test from the agricolae package in R (de Mendiburu, 2019).

### Extraction and analysis of anthocyanins and flavonoids

Anthocyanins were quantified from seedling extracts as described previously (Yin et al., 2012). Approximately 50 mg fresh weight of seedlings were ground and pigments were extracted in 250 μl methanol with 1% [v/v] HCl at 4°C for at least 1 h. Clear supernatants were collected, and absorbance was measured at 530 nm and 655 nm. The amount of anthocyanins was calculated as (A530 - 0.25 * A655) / m, where m is the fresh weight of the seedlings.

Flavonol profiles were analyzed by high-performance thin layer chromatography (HPTLC) as previously described (Stracke et al., 2010). In brief, 50 mg of seedlings were harvested, ground, and incubated with 100 μl of 80% [v/v] methanol on a shaker for 10 min at 70°C before centrifugation. Clear supernatants were then collected and 10 μl of samples was spotted on silica HPTLC plates using capillary tubes. The methanolic extracts were then separated in a mobile phase consisting of a mixture of 5 ml ethyl acetate, 600 μl formic acid, 600 μl acetic acid glacial, and 1.3 ml water. After migration, the plate was dried and the flavonol staining was revealed under a 365-nm UV lamp after spraying the chromatogram with a 1% [w/v] diphenylboric acid 2-aminoethylester (DPBA; Roth) solution in 80% [v/v] methanol.

### Immunoblot analysis

For analysis of protein levels by immunoblotting, proteins were extracted from seedlings using an extraction buffer consisting of 50 mM Na-phosphate pH 7.4, 150 mM NaCl, 10% [v/v] glycerol, 5 mM EDTA, 0.1% [v/v] Triton X-100, 1 mM DTT, 2 mM Na3VO4, 2 mM NaF, 1% [v/v] Protease Inhibitor Cocktail (Sigma), and 50 μM MG132 (Arongaus et al., 2018).

To determine the dimer/monomer status of UVR8 by SDS-PAGE, proteins were extracted in an extraction buffer composed of 150 mM NaCl, 50 mM Tris-HCl pH 7.6, 2 mM EDTA, 1% [v/v] Igepal (Sigma), 1% [v/v] Protease Inhibitor Cocktail (Sigma), 10 μM MG132, and 10 μM ALLN (VWR) (Heijde and Ulm, 2013). Extracts were then loaded on SDS-PAGE gels without prior heat denaturation and gels were exposed to broadband UV-B for 30 min before transfer, as described previously (Rizzini et al., 2011).

For HY5 immunoblots, the following extraction buffer was used: 50 μM EDTA, 0.1 M Tris-HCl pH 8, 0.7% [w/v] SDS, 10 mM NaF, cOmplete^TM^ EDTA-free Protease Inhibitor Cocktail Tablet (Roche), 1 mM DTT, 0.25 M NaCl, 15 mM β-glycerolphosphate, and 15 mM p-nitrophenyl phosphate (Oravecz et al., 2006).

Following electrophoretic separation in SDS-PAGE gels, proteins were transferred to PVDF membranes (Roth) according to the manufacturer’s instructions (iBlot dry blotting system, Thermo Fisher Scientific); however, to analyze RUP2 levels, proteins were liquid-transferred to nitrocellulose membranes (Bio-Rad). Membranes were blocked in 10% [w/v] milk, except for HY5 immunoblots, which were blocked by drying the membrane.

anti-UVR8^(426–440)^ (Favory et al., 2009), anti-UVR8^(1–15)^ (Yin et al., 2015), anti-GFP (Living Colors® A.v. Monoclonal Antibody, JL-8; Clontech), anti-CHS (sc-12620; Santa Cruz Biotechnology), anti-UGPase (AS05086, Agrisera), anti-actin (A0480; Sigma-Aldrich), anti-HY5 (Oravecz et al., 2006), and anti-RUP2 (Arongaus et al., 2018) were used as primary antibodies, and corresponding horseradish peroxidase-conjugated anti-rabbit, anti-mouse, and anti-goat (Dako) immunoglobulins were used as secondary antibodies. Signal detection was done on an Amersham Imager 680 camera system (GE Healthcare) using the ECL Select Western Blotting Detection Reagent (GE Healthcare).

### Protein crosslinking

Crosslinking experiments were performed as described before (Rizzini et al., 2011). Proteins were extracted from seedlings in PBS containing 0.1% [v/v] Igepal, 1 mM PMSF, 10 mM leupeptine, 1% [v/v] protease inhibitor cocktail for plants (Sigma), 10 μM MG132, and 10 μM ALLN (VWR). Extracts were then centrifuged and clear supernatants were incubated with 2 mM dithiobis(succinimidyl propionate) (DSP; ThermoFisher) for 30 min at 4°C on a rotary shaker. Crosslinking was then quenched with 50 mM Tris pH 7.6 for 15 min at room temperature. Samples were then heat denatured in SDS-PAGE loading buffer without reducing agent. To reverse crosslinking, 5% [v/v] β-mercaptoethanol was added prior to heat-based denaturation.

### Yeast two-hybrid (Y2H) analysis

To test the UVR8–COP1 interaction, *COP1* was inserted into pGADT7-GW (Marrocco et al., 2006; Heijde et al., 2013) and *UVR8*, *UVR8^G101S^*, *UVR8^W285A^, UVR8^G101S,W285A^, UVR8^W285F^, UVR8^G101S,W285F^*, and *UVR8^D96N,D107N^* were cloned into pBTM116-D9-GW (Stelzl et al., 2005; Heijde et al., 2013). The L40 yeast stain (Vojtek and Hollenberg, 1995) was used for transformation using the lithium acetate–based transformation protocol (Gietz, 2014).

To test the interaction of UVR8 with RUP2, *RUP2* was cloned into pGBKT7-GW (Lu et al., 2010) and transformed into the Y2H Gold strain (Clontech). *UVR8* and *UVR8^G101S^* were introduced into pGADT7-GW (Marrocco et al., 2006) and transformed into the Y187 strain (Wade Harper et al., 1993). The relevant pairs were then combined by mating.

Transformed yeast cells were selected on SD/-Trp/-Leu medium (Foremedium) for non-selective growth and on a SD/-Trp/-Leu/-His medium for selective growth. To quantify the interactions using the LacZ reporter, yeast strains were grown for 2 d on non-selective medium and the β-galactosidase enzymatic activity was determined in an assay using red-β-D-galactopyranoside (CPRG, Roche Applied Science) as substrate (Yeast Protocols Handbook, Clontech). For UV-B treatments, yeast cells were irradiated with 1.5 μmol m^-2^ s^-1^ of narrow-band UV-B provided by Philips TL20W/01RS tubes.

### Protein purification from Sf9 cell cultures

*Spodoptera frugiperda* Sf9 cells (Thermofisher) were cultured in Sf-4 Baculo Express insect cell medium (Bioconcept, Switzerland). Each of the recombinant COP1^349–675^, UVR8 full-length, and UVR8^12–381^ proteins were produced as described before (Lau et al., 2019). The desired Arabidopsis full-length or truncated coding sequence was PCR amplified or NcoI/NotI digested from codon-optimized genes (Geneart) for expression in Sf9 cells. Mutant UVR8 constructs were produced using an enhanced plasmid mutagenesis protocol (Liu and Naismith, 2008). All were cloned into a modified pFastBac (Geneva Biotech) insect cell expression vector via NcoI/NotI restriction enzyme sites or by Gibson assembly (Gibson et al., 2009). The modified pFastBac vector contains a tandem N-terminal His10-Twin-Strep-tags followed by a TEV (tobacco etch virus protease) cleavage site. pFastBac constructs were transformed into DH10MultiBac cells (Geneva Biotech), following which white colonies indicating successful recombination were selected and bacmids were purified by the alkaline lysis method. Sf9 cells were transfected with the desired bacmid with Profectin (AB Vector). eYFP-positive cells were observed after 1 week and subjected to one round of viral amplification. Amplified, untitred P2 virus (between 5–10% culture volume) was used to infect Sf9 cells at a density between 1–2 x 10^6^ cells/ml. Cells were incubated for 72 h at 28°C before the cell pellet was harvested by centrifugation at 2000 × *g* for 20 min and stored at –20°C.

Pellets from every liter of Sf9 cell culture were dissolved in 25 ml of buffer A (300 mM NaCl, 20 mM HEPES 7.4, 2 mM β-ME), supplemented with 10% [v/v] glycerol, 5 μl Turbonuclease, and 1 Roche cOmplete^TM^ protease inhibitor tablet. Dissolved pellets were lysed by sonication and insoluble materials were separated by centrifugation at 60,000 × *g* for 1 h at 4°C. The supernatant was filtered through tandem 1-μm and 0.45-μm filters before Ni^2+^-affinity purification (HisTrap excel, GE Healthcare). Ni^2+^-bound proteins were washed with buffer A and eluted directly onto a coupled Strep-Tactin Superflow XT column (IBA) by buffer B (500 mM NaCl, 500 mM imidazole pH 7.4, 20 mM HEPES pH 7.4). Tandem-Strep-tagged-bound proteins on the Strep-Tactin column were washed with buffer A and eluted with 1x Buffer BXT (IBA). Proteins were cleaved overnight at 4°C with TEV protease. Cleaved proteins were subsequently purified from the protease and affinity tag by a second Ni^2+^-affinity column or by gel filtration on a Superdex 200 Increase 10/300 GL column (GE Healthcare). Proteins were concentrated to 3–10 mg/ml and either used immediately or aliquoted and quickly frozen at –80°C. Typical purifications were from pellets of 2–5 liters of insect cell culture. All protein concentrations were measured by absorption at 280 nm and calculated from their molar extinction coefficients. Molecular weights of all proteins were confirmed by MALDI-TOF mass spectrometry. SDS-PAGE gels to assess protein purity are shown in Supplementary Fig. 14. For UVR8 monomerization and activation by UV-B, purified UVR8 proteins were exposed for 60 min at max intensity (69 mA) under UV-B LEDs (Roithner Lasertechnik GmbH) on ice.

### Analytical size-exclusion chromatography

Gel filtration experiments were performed using a Superdex 200 Increase 10/300 GL column (GE Healthcare) pre-equilibrated in 150 mM NaCl, 20 mM HEPES 7.4, and 2 mM β-ME. 500 μl of the respective protein solution (∼4 μM per protein) was loaded sequentially onto the column and elution at 0.75 ml/min was monitored by UV absorbance at 280 nm.

### *In vitro* methylation

150 μl of 4 mg/ml COP1/UVR8^12–381, D96N,D107N^ complex was diluted to 500 μl using buffer (150 mM NaCl, 20 mM HEPES pH 7.4). 20 μl of 1 M dimethylaminoborane (DMAB) and 40 μl of formaldehyde was added to the protein mixture at 4°C and left rotating for 2 h. The addition of DMAB and formaldehyde was repeated once. 10 μl of 1 M DMAB was subsequently added and the mixture was left on ice overnight. The reaction was quenched with 125 μl of 1 M Tris pH 8, concentrated to 500 μl and loaded onto a Superdex 200 Increase 10/300 GL column. Methylation was confirmed by MALDI-TOF mass spectrometry with ∼13 free amines methylated on UVR8^D96N,D107N^.

### Protein crystallization and data collection

Crystals of truncated UVR8 (residues 12–381) mutants were grown in sitting drops and appeared after several days (UVR8^D96N,D107N^, UVR8^D96N,D107N,W285A^) to 1 year (UVR8^G101S,W285A^) at 20°C in drops where 5 mg/ml of UVR8^D96N,D107N,W285A^ or UVR8^G101S,W285A^ was mixed in a protein:buffer ratio of 1:1. UVR8^D96N,D107N,W285A^ crystals formed in 0.2 M NaNO3, 22% [w/v] PEG 3,350, whereas UVR8 UVR8^G101S,W285A^ crystals formed in 0.1 M Bis-Tris propane pH 8.5, 0.2 M NaNO3, 20% [w/v] PEG 3,350. UVR8^D96N,D107N^ crystals formed when *in vitro* methylated COP1/UVR8^D96N,D107N^ complex at 1.8 mg/ml was mixed in a ratio of 2:1 protein:buffer in 0.1 M Tris pH 8.5, 0.1 M NaCl, 30% [w/v] PEG 4,000. Crystals were harvested and cryoprotected in mother liquor supplemented with 25% [v/v] glycerol for UVR8^D96N,D107N^ and UVR8^G101S,W285A^ or with 20% [w/v] PEG 400 for UVR8^D96N,D107N,W285A^ and frozen in liquid nitrogen.

Native datasets were collected at beam line PX-III of the Swiss Light Source (Villigen) with λ=1.03 Å. All datasets were processed with XDS (Kabsch, 1993) and scaled with AIMLESS as implemented in the CCP4 suite (Winn et al., 2011).

### Crystallographic structure solution and refinement

The structures of the mutant UVR8 versions were solved by molecular replacement as implemented in the program Phaser (McCoy et al., 2007), using PDB-ID 4D9S as the initial search model. The final structures were determined after iterative rounds of model-building in COOT (Emsley and Cowtan, 2004), followed by refinement in REFMAC5 (Murshudov et al., 2011) and phenix.refine (Adams et al., 2010). Final statistics were generated using phenix.table_one. Structural diagrams were rendered in UCSF Chimera (Pettersen et al., 2004) and UCSF ChimeraX (Goddard et al., 2018).

### Grating-coupled interferometry (GCI)

The Creoptix WAVE system (Creoptix AG), a label-free surface biosensor, was used to perform GCI experiments. All experiments were performed on 2PCH or 4PCH WAVEchips (quasi-planar polycarboxylate surface; Creoptix AG). After a borate buffer conditioning (100 mM sodium borate pH 9.0, 1 M NaCl; Xantec) COP1 (ligand) was immobilized on the chip surface using standard amine-coupling: 7-min activation (1:1 mix of 400 mM *N*-(3-dimethylaminopropyl)-*N*′-ethylcarbodiimide hydrochloride and 100 mM *N*-hydroxysuccinimide (both Xantec)), injection of COP1 (10 μg/ml) in 10 mM sodium acetate pH 5.0 (Sigma) until the desired density was reached, and final quenching with 1 M ethanolamine pH 8.0 for 7 min (Xantec). For a typical experiment, the analyte (UVR8) was injected in a 1:3 dilution series in 150 mM NaCl, 20 mM HEPES 7.4, 2 mM β-ME at 25°C. Blank injections were used for double referencing and a dimethylsulfoxide (DMSO) calibration curve for bulk correction. Analysis and correction of the obtained data was performed using the Creoptix WAVEcontrol software (applied corrections: X and Y offset; DMSO calibration; double referencing) and a one-to-one binding model or a heterogenous ligand model with bulk correction was used to fit all experiments. Data of GCI binding assays are reported with errors as indicated in their figure legends.

### Data Availability

The atomic coordinates of complexes have been deposited with the following Protein Data Bank (PDB) accession codes: UVR8^D96N,D107N^: **6XZL** (https://www.rcsb.org/structure/6XZL), UVR8^D96N,D107N,W285A^: **6XZM** (https://www.rcsb.org/structure/6XZM), UVR8^G101S,W285A^: **6XZN** (https://www.rcsb.org/structure/6XZN).

## ACKNOWLEDGEMENTS

We thank Richard Chappuis for excellent technical assistance and José Manuel Nunes (BioSC, Geneva) for helpful discussions regarding statistical analyses. This work was supported by the University of Geneva, an HHMI International Research Scholar Award to M.H., the Swiss National Science Foundation (grant number 31003A_175774 to R.U.), and the European Research Council (ERC) under the European Union’s Seventh Framework Programme (grant no. 310539 to R.U.). R.P. was supported by an iGE3 PhD Salary Award, K.L. was supported by an EMBO Long-term Fellowship (ALTF 493-2015).

---

**Supplementary Fig. 1.**
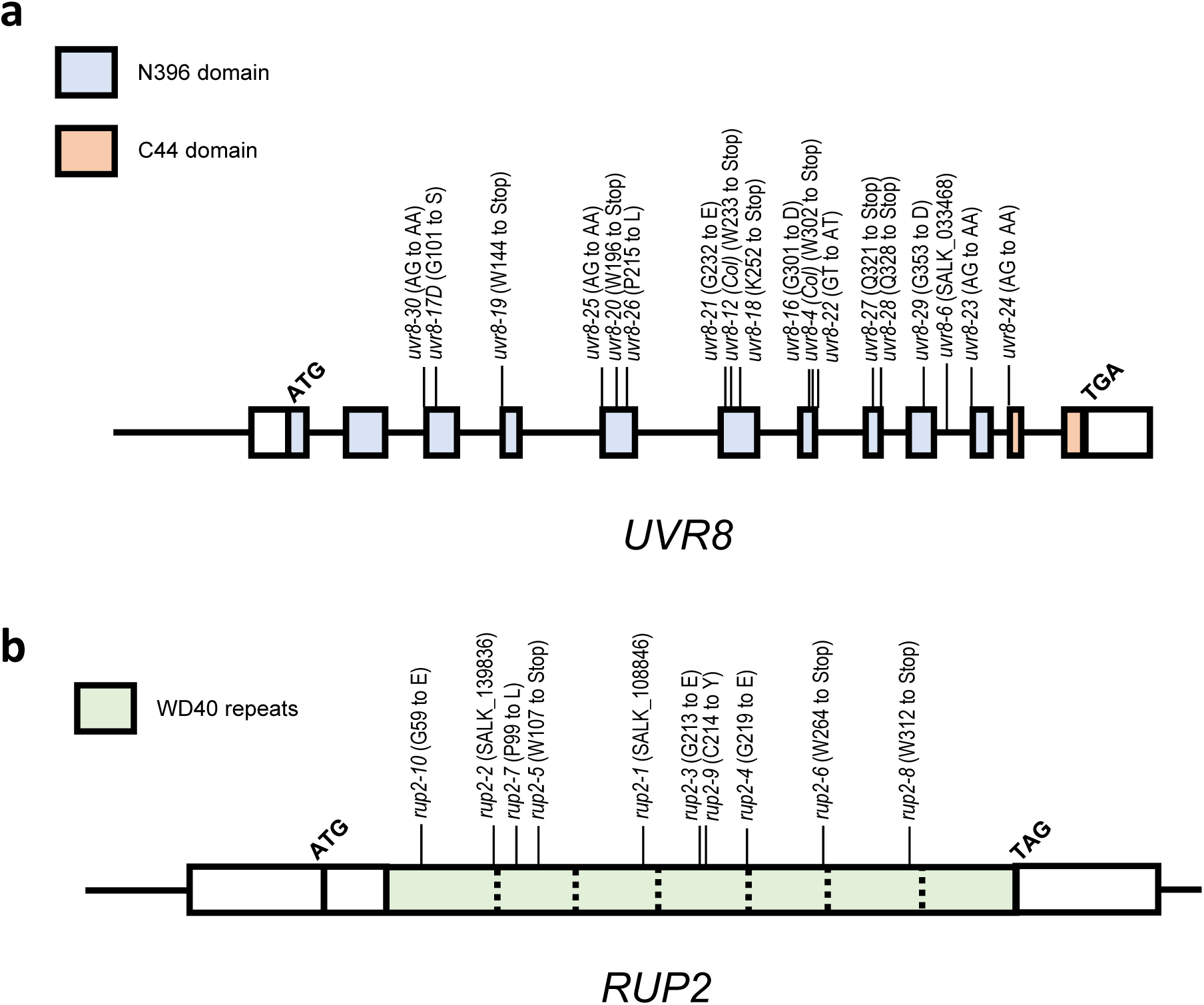
Novel *uvr8* and *rup2* loss-of-function alleles. **a** Structure of the *UVR8* gene and new mutant alleles. Boxes indicate exons. The sequence corresponding to the N396 domain is colored in blue, the C44 domain is in orange (Yin et al., 2015). Novel alleles identified in the hypocotyl length–based screen are indicated (Col accession), in addition to the previously characterized *uvr8-6* T-DNA insertion mutant (Favory et al., 2009), and already known alleles previously described in other accessions: *uvr8-4* (originally reported in L*er*) (Cloix et al., 2012) and *uvr8-12* (originally reported in Ws) (Favory et al., 2009). Additionally *uvr8-17D* is indicated. **b** Structure of the *RUP2* gene and new mutant alleles. Boxes indicate exons. The sequence colored in green represents the seven WD40 repeats (separated by dashed lines). The T-DNA insertion lines *rup2-1* (Gruber et al., 2010) and *rup2-2* (Arongaus et al., 2018) are indicated, as well as all novel alleles.

**Supplementary Fig. 2.**
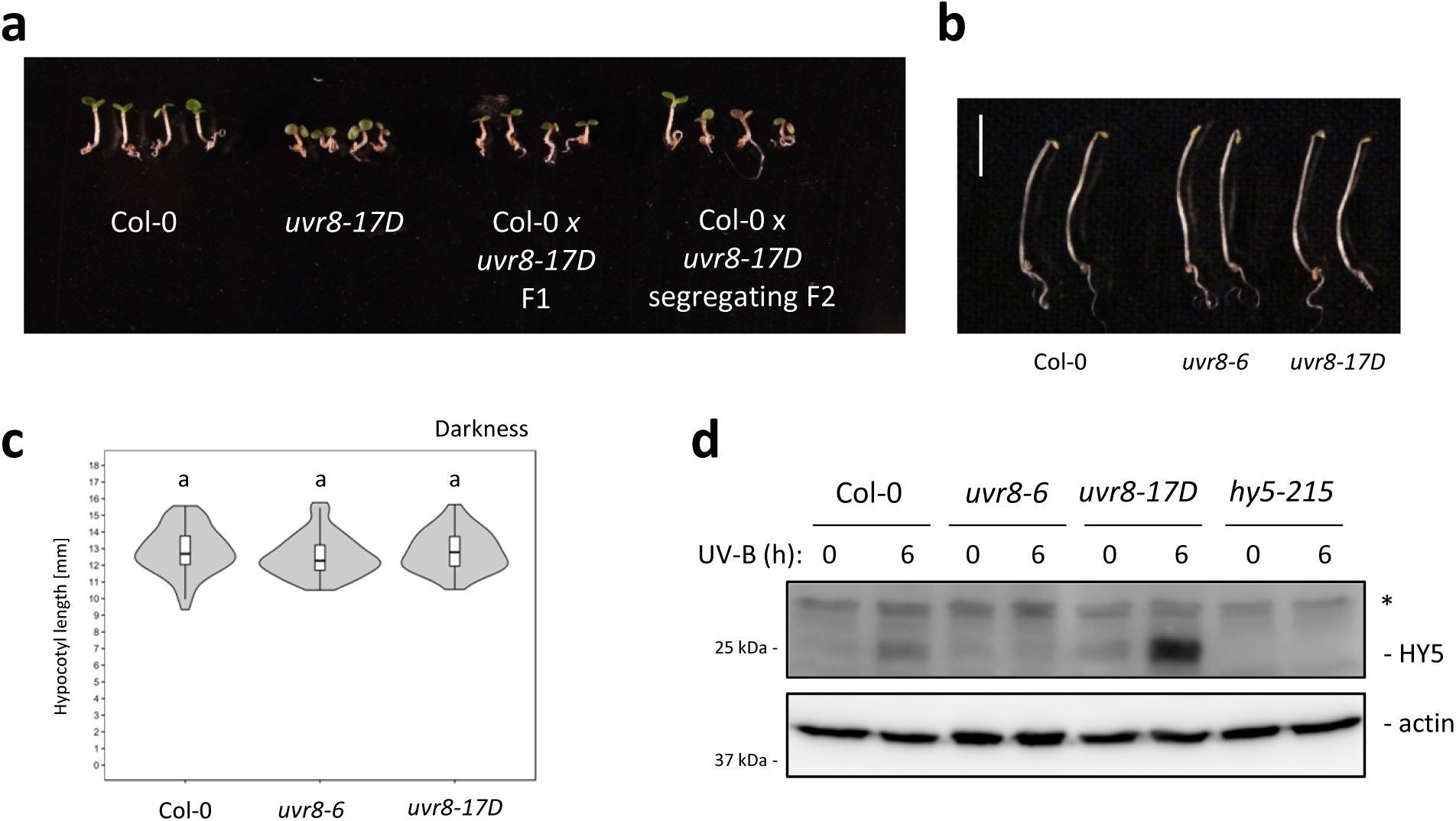
Characterization of the *uvr8-17D* allele. **a** Representative images of the seedling phenotypes of wild type (Col-0), *uvr8-17D*, and F1 as well as segregating F2 progeny of a Col-0 x *uvr8-17D* cross, following seedling growth in white light supplemented with UV-B. **b** Representative images of wild-type, *uvr8-6*, and *uvr8-17D* seedlings grown in darkness. Bar = 5 mm. **c** Quantification of hypocotyl length of 4-d-old seedlings grown in darkness, representing genotypes described in **b** (*N* > 60). Shared letters indicate no statistically significant difference in the means (*P* < 0.05). **d** Immunoblot analysis of HY5 and actin (loading control) protein levels in wild-type, *uvr8-6*, *uvr8-17D*, and *hy5-215* seedlings grown in white light for 4 d, and either exposed or not exposed to supplemental UV-B for 6 h. Asterisk indicates nonspecific cross-reacting bands.

**Supplementary Fig. 3.**
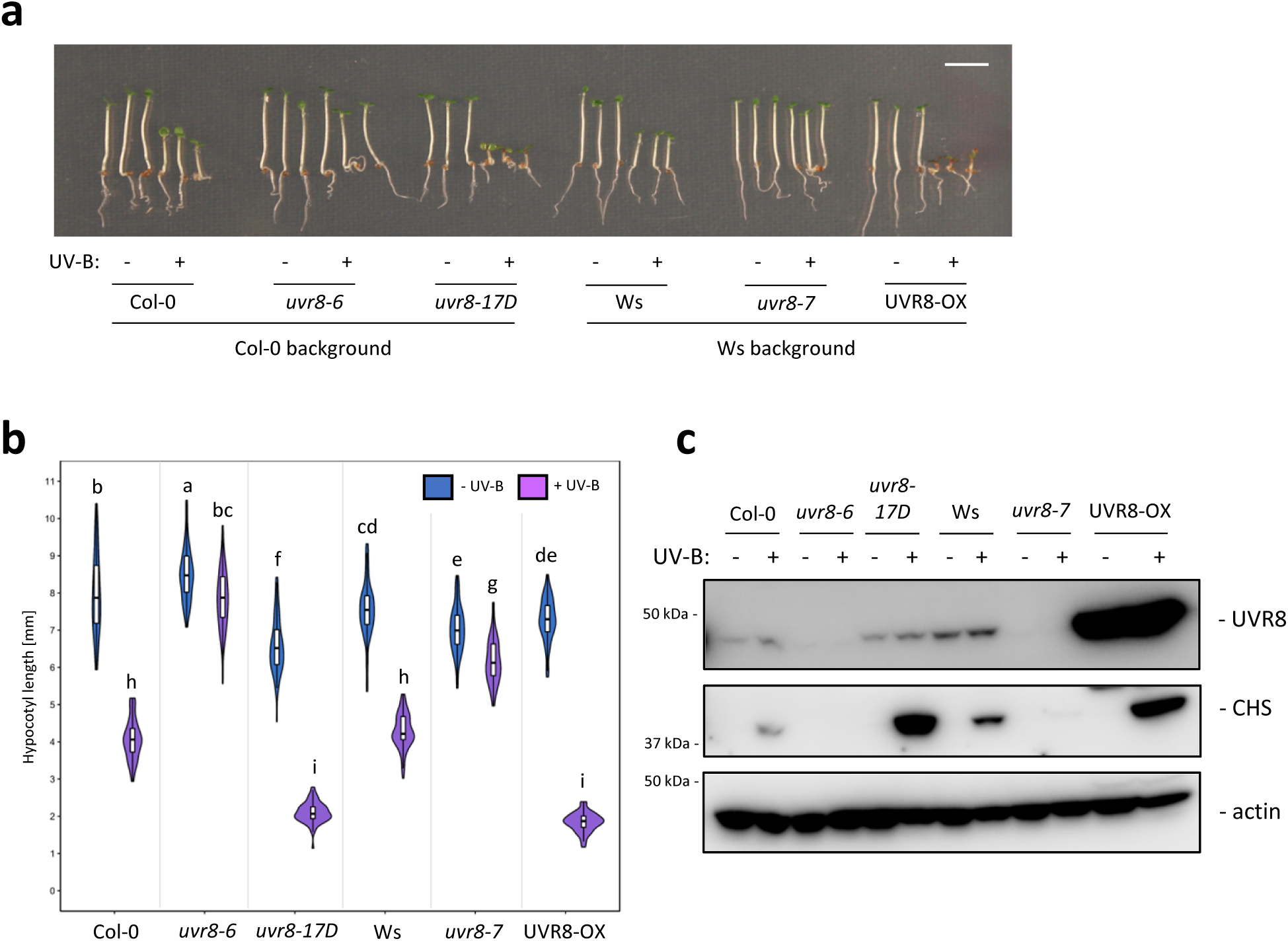
Comparison of *uvr8-17D* and UVR8-OX. **a** Representative images of wild-type (Col-0), *uvr8-6*, *uvr8-17D*, wild-type (Ws), *uvr8-7*, and *uvr8-7*/Pro_35S_:UVR8 (UVR8-OX) seedlings grown for 4 d in white light or white light supplemented with UV-B. Bar = 5 mm. **b** Quantification of hypocotyl length of seedlings described in **a** (*N* > 60). Shared letters indicate no statistically significant difference in the means (*P* < 0.05). **c** Immunoblot analysis of UVR8, CHS, and actin (loading control) protein levels in seedlings grown as described in **a**.

**Supplementary Fig. 4.**
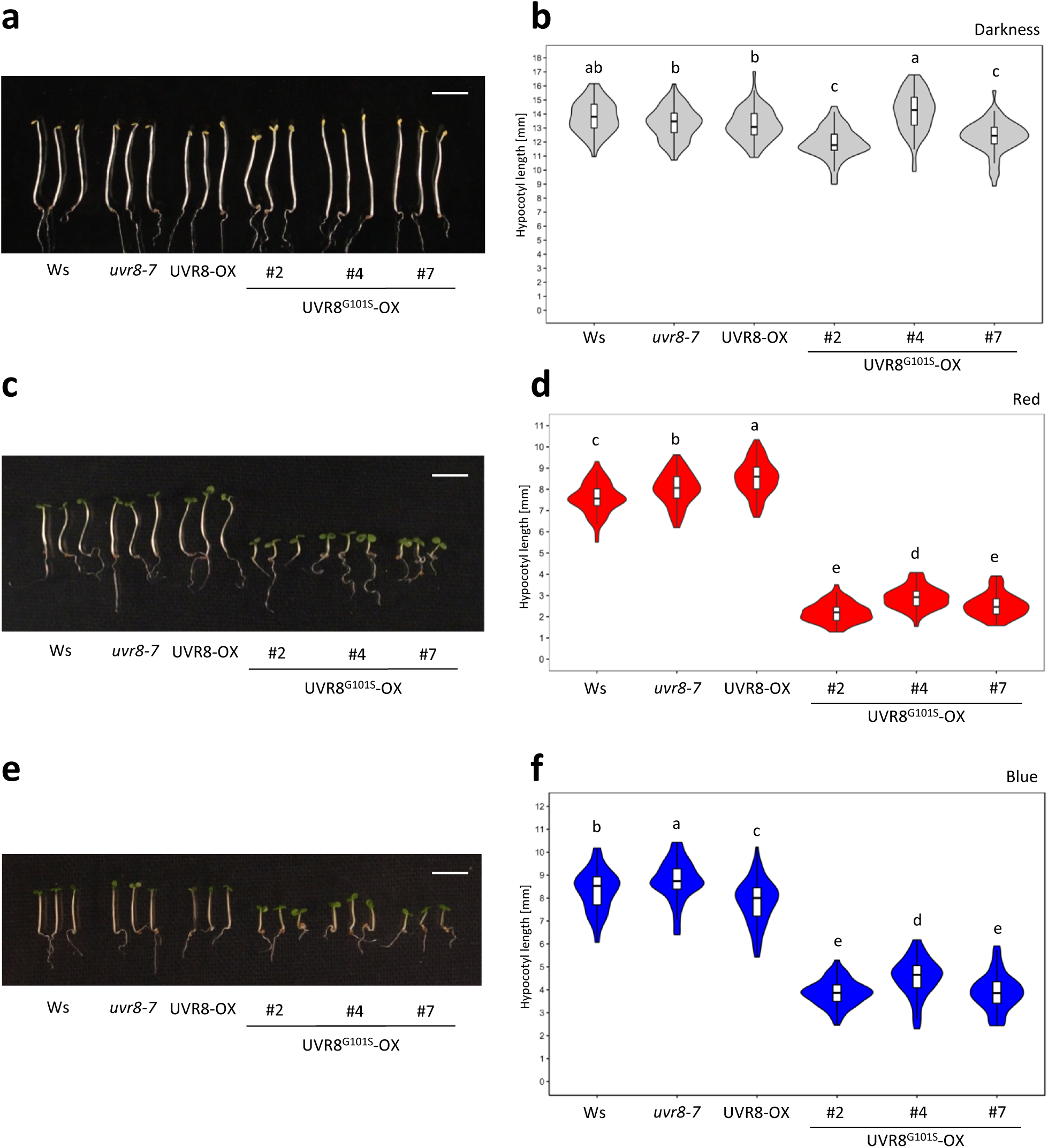
UVR8^G101S^ overexpression lines show weak constitutive photomorphogenesis. **a,c,e** Representative images of seedlings of wild type (Ws), *uvr8-7*, *uvr8-7*/Pro_35S_:UVR8 (UVR8-OX), and three independent *uvr8-7*/Pro_35S_:UVR8^G101S^ (UVR8^G101S^-OX #2, #4, and #7) lines grown **a** in darkness, **c** under 30 μmol m^-2^ s^-1^ of red light, or **e** under 5 μmol m^-2^ s^-1^ of blue light. Bar = 5 mm. **b**,**d**,**f** Quantification of hypocotyl length of seedlings described in **a,c,e**, respectively (*N* > 60). Shared letters indicate no statistically significant difference in the means (*P* < 0.05).

**Supplementary Fig. 5.**
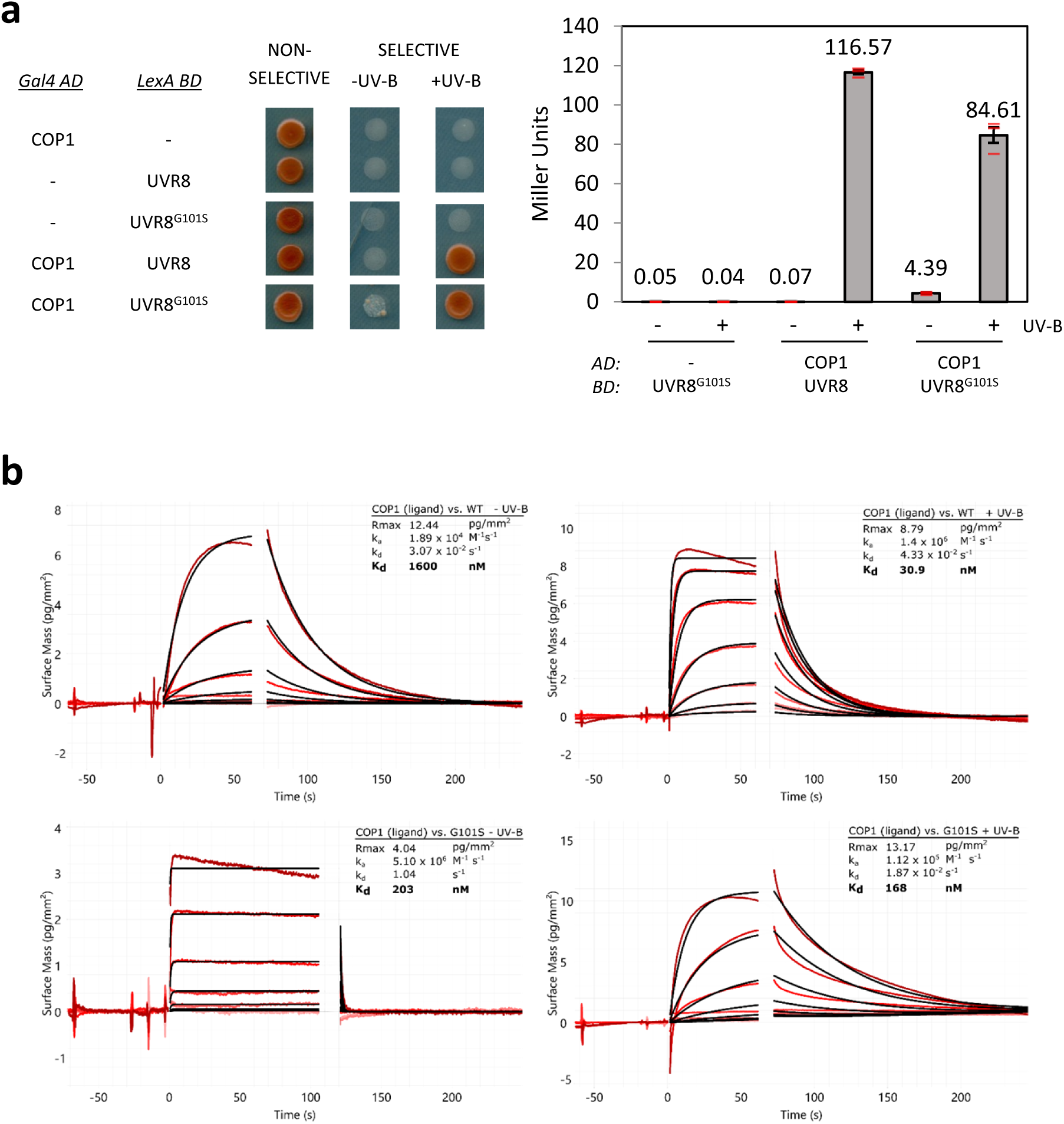
UVR8^G101S^ interacts with COP1 in a UV-B-dependent manner. **a** Y2H analyses of the interactions between COP1 and UVR8/UVR8^G101S^ in the presence or absence of UV-B. Left: growth assay on selective SD/-Trp/-Leu/-His medium. Right: quantitative β-galactosidase assay. AD, activation domain; BD, DNA binding domain. **b** Binding kinetics of full-length UVR8 and UVR8^G101S^ versus the COP1 WD40 domain obtained by GCI experiments. Sensorgrams of protein injected are shown in red, with their respective heterogenous ligand binding model fits in black. The following amounts were typically used: ligand: COP1 (2,000 pg/mm^2^); analyte: UVR8 (2 μM highest concentration). ka = association rate constant, kd = dissociation rate constant, Kd = dissociation constant.

**Supplementary Fig. 6.**
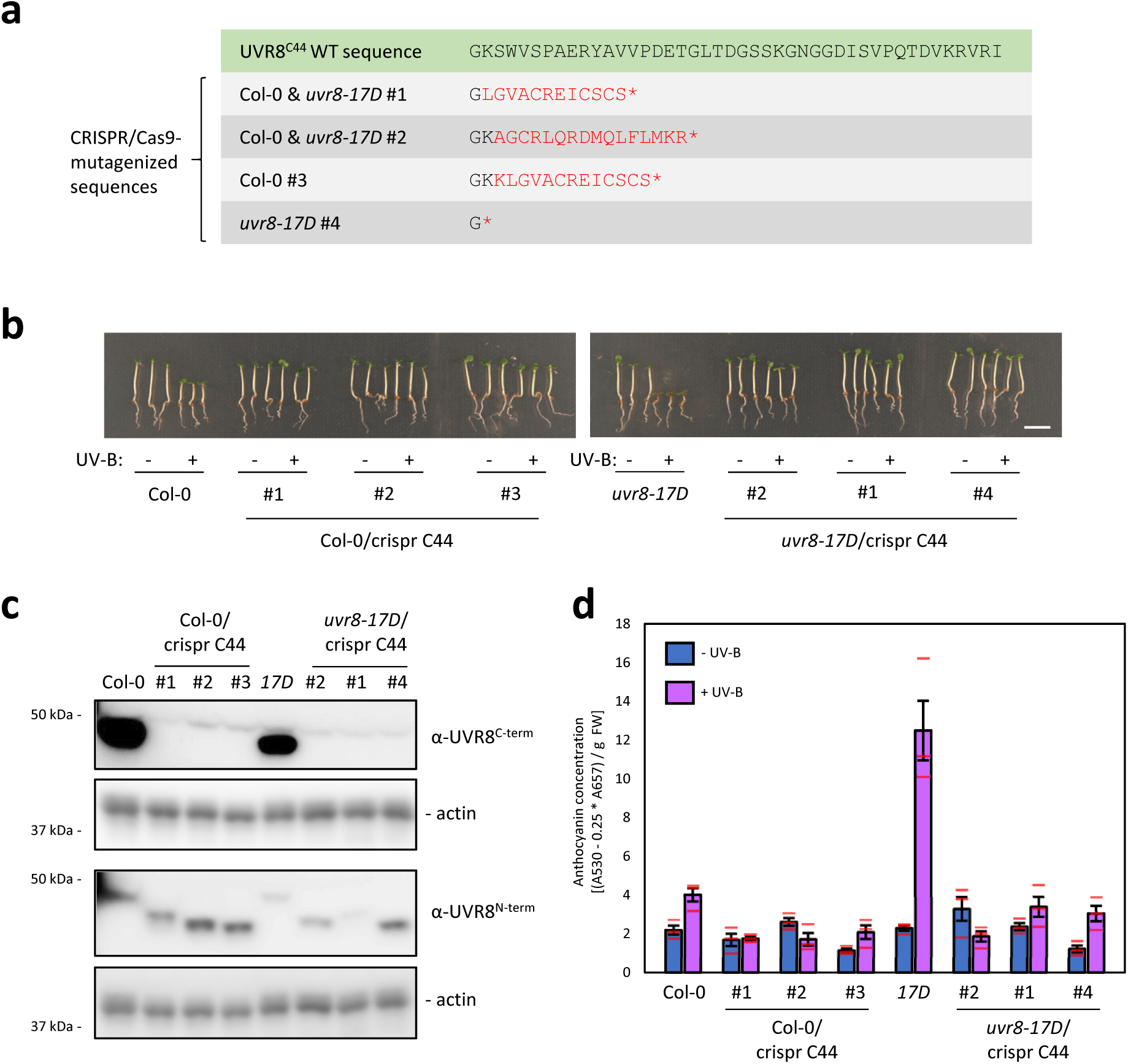
Deletion of the VP motif–containing C-terminus of UVR8 abolishes the UV-B response in *uvr8-17D*. **a** Mutations induced by CRISPR/Cas9 directed against the *UVR8* sequence encoding the UVR8 C-terminus. The UVR8^C44^ wild-type sequence is shown, as well as the mutated sequences documented in Col-0 and/or *uvr8-17D* backgrounds. Residues generated as a consequence of frameshift are indicated in red, * indicates a newly formed translation stop codon. **b** Representative images of wild-type (Col-0) and *uvr8-17D* seedlings alongside three respective independent mutant lines containing CRISPR/Cas9-generated C-terminal C44 truncations (Col-0/crispr C44 #1–3, and *uvr8-17D*/crispr C44 #2, #1, and #4) grown in white light or white light supplemented with UV-B. Bar = 5 mm. **c** Immunoblot analysis of UVR8 and actin (loading control) protein levels in the lines described in **b** (*uvr8-17D* = *17D*). For analysis of UVR8 levels, antibodies specifically recognizing the N-terminus (α-UVR8^N-term^ = α-UVR8^1-15^) or the C-terminus (α-UVR8^C-term^ = α-UVR8^426-440^) of UVR8 were used. **d** Anthocyanin concentration in the lines described in **b**; values of independent measurements (red bars), means, and SEM are shown (*N* = 3).

**Supplementary Fig. 7.**
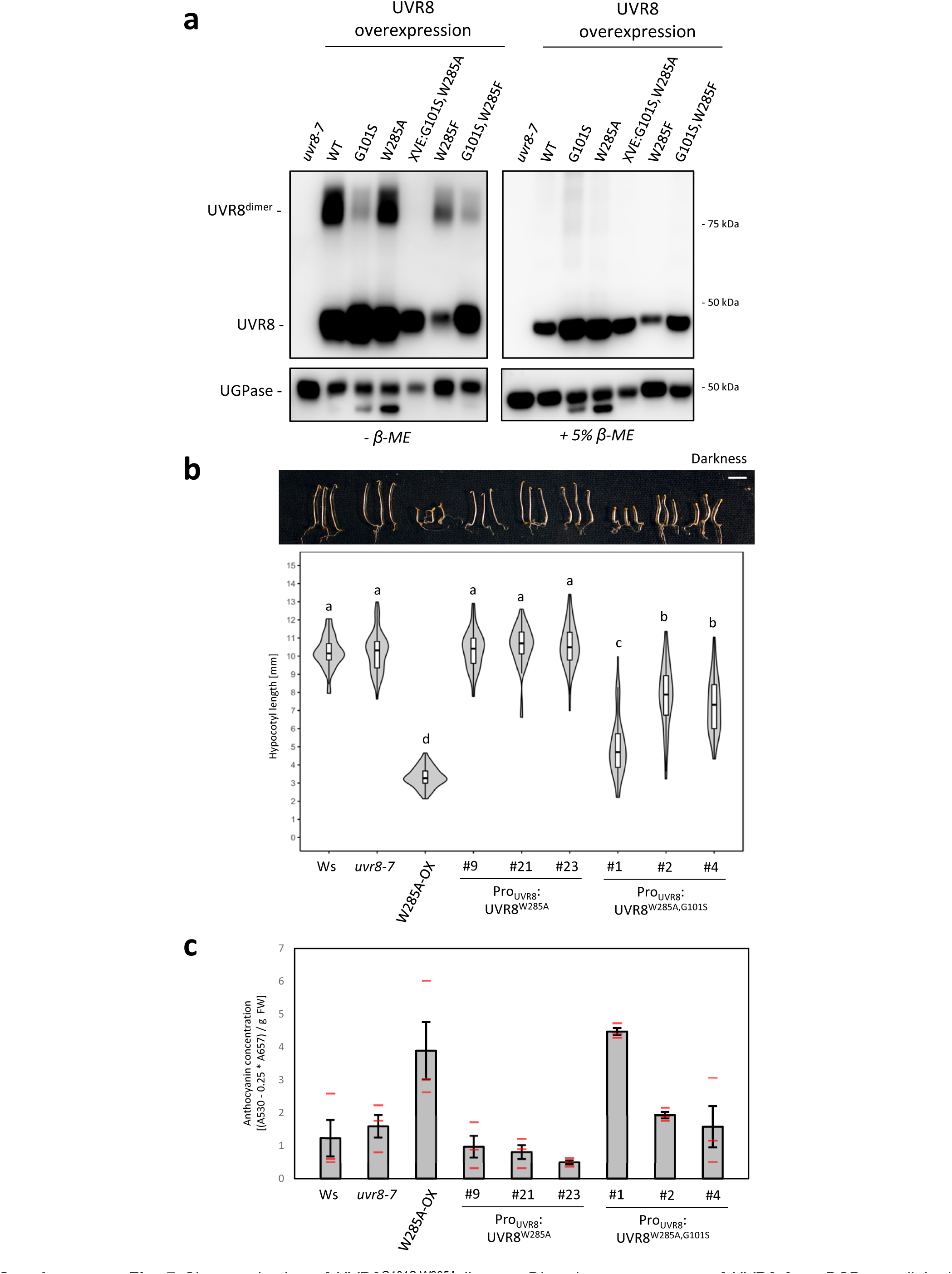
Characterization of UVR8^G101S,W285A^ lines. **a** Dimer/monomer status of UVR8 from DSP-crosslinked extracts of various UVR8-overexpressing lines under a 35S or XVE-responsive promoter. UVR8^W285A^- and XVE:UVR8^G101S,W285A^-expressing lines were grown on 5 μM estradiol. Crosslinking was reversed by addition of 5% β-mercaptoethanol. UGPase is shown as loading control. **b** Representative images and quantification of hypocotyl length of seedlings of wild type (Ws), *uvr8-7*, *uvr8-7*/Pro_35S_:UVR8^W285A^ (W285A-OX), and three independent lines each of *uvr8-7*/Pro_UVR8_:UVR8^W285A^ (#9, #21, and #23) and *uvr8-7*/Pro_UVR8_:UVR8^G101S,W285A^ (#1, #2, and #4) grown in darkness (*N* > 60). Shared letters indicate no statistically significant difference in the means (*P* < 0.05). Bar = 5 mm. **c** Anthocyanin concentration in the lines described in **b**. Values of independent measurements (red bars), means, and SEM are shown (*N* = 3).

**Supplementary Fig. 8.**
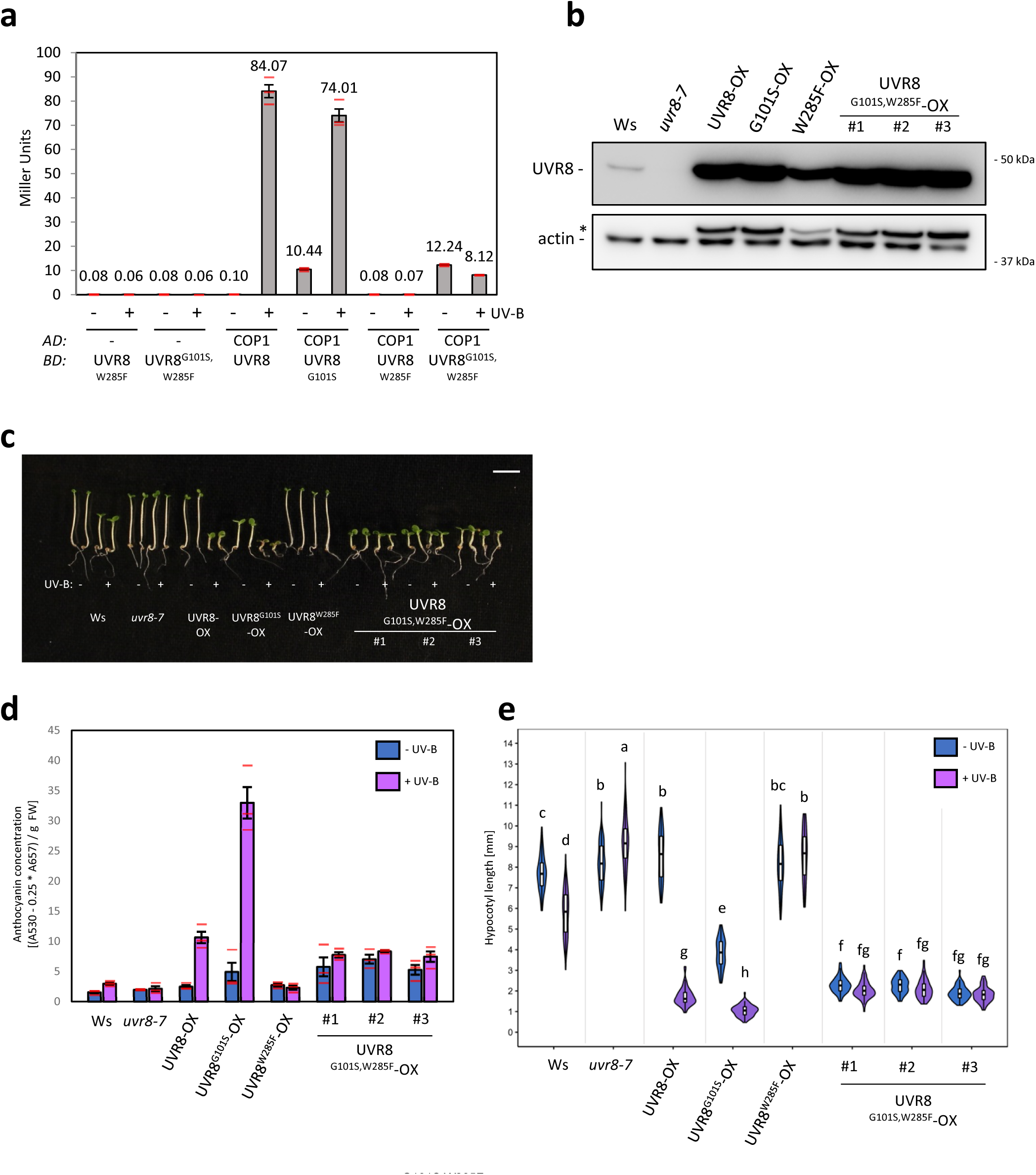
Overexpression of UVR8^G101S,W285F^ results in weak constitutive photomorphogenesis similar to non-UV-B exposed UVR8^G101S^. **a** Quantitative Y2H analysis of the interaction between COP1 and UVR8, UVR8^G101S^, UVR8^W285F^, and UVR8^G101S,W285F^ in the absence of UV-B. AD, activation domain; BD, DNA binding domain. **b** Immunoblot analysis of UVR8 and actin (loading control) protein levels in wild type (Ws), *uvr8-7*, *uvr8-7*/Pro_35S_:UVR8 (UVR8-OX), *uvr8-7*/Pro_35S_:UVR8^G101S^ #2 (G101S-OX), *uvr8-7*/Pro_35S_:UVR8^W285F^ (W285F-OX), and three independent lines of *uvr8-7*/Pro_35S_:UVR8^G101S,W285F^ (#1–3). Asterisk indicates residual UVR8 signal after stripping of the PVDF membrane. **c** Representative images of seedlings described in **b** grown for 4 d in white light or white light supplemented with UV-B. Bar = 5 mm. **d** Anthocyanin concentration in seedlings described in **c**. Values of independent measurements (red bars), means, and SEM are shown (*N* = 3). **e** Quantification of hypocotyl length in seedlings described in **c** (*N* > 60). Shared letters indicate no statistically significant difference in the means (*P* < 0.05).

**Supplementary Fig. 9.**
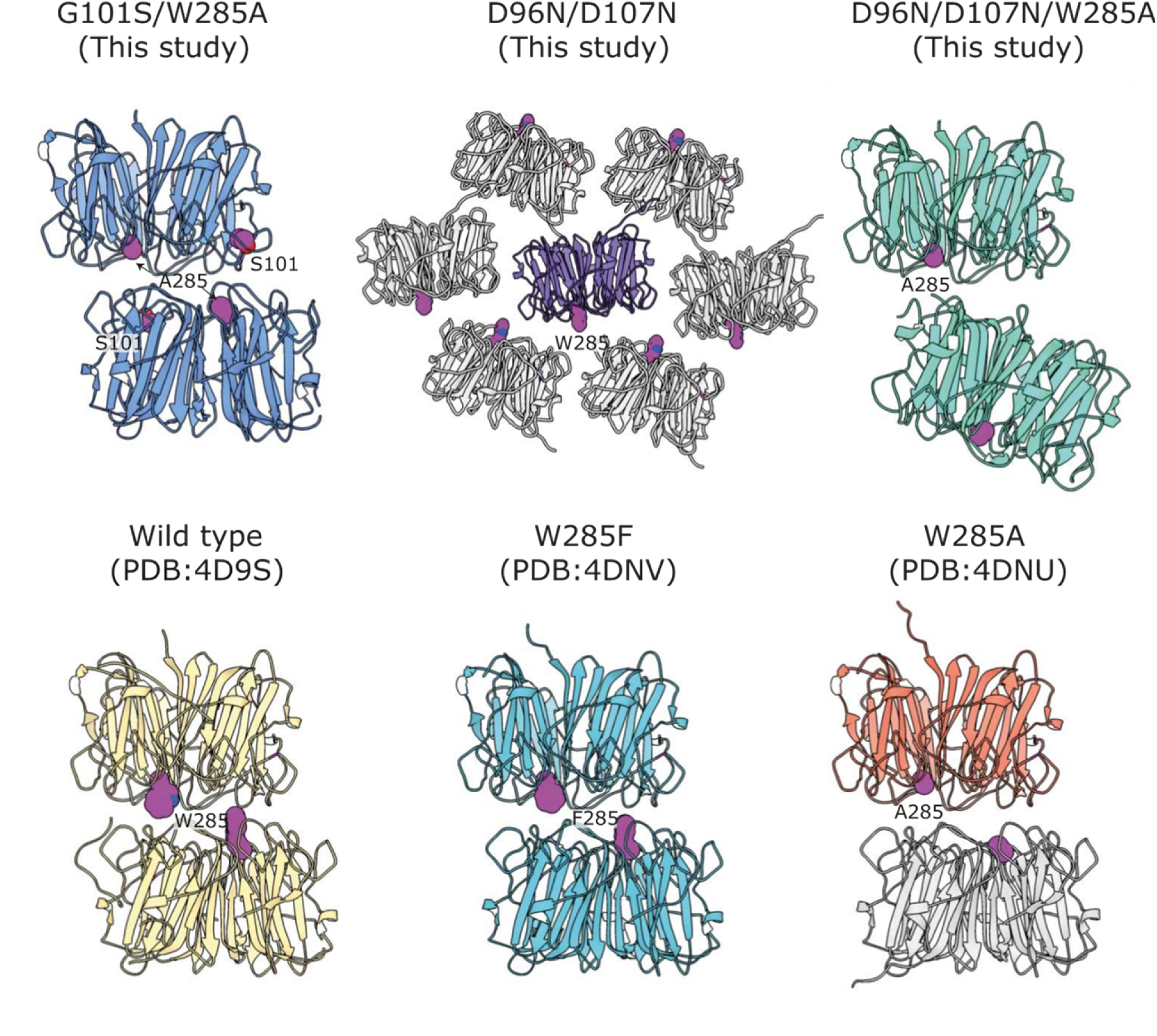
Lattice interactions in different UVR8 variant crystal forms. UVR8 variants are depicted with colors and each represents the UVR8 present in one unit cell. Crystallographic symmetry partners are generated and depicted in gray to show higher-order assemblies when necessary. The residues at position 285 or 101 are highlighted in magenta spheres for orientation. UVR8^G101S,W285A^ (G101S/W285A), UVR8 (wild type), and UVR8^W285F^ (W285F) crystallize as conventional ‘wild-type’ (top-to-top) symmetric dimers. UVR8^W285A^ (W285A) crystallizes with one molecule in the asymmetric unit, but form a canonical UVR8 dimer by symmetry within the crystal lattice. UVR8^D96N,D107N^ (D96N/D107N) crystallizes also as a monomer and its symmetry mates show various conformations that do not correspond to a dimer. UVR8^D96N,D107N,W285A^ (D96N/D107N/W285A) crystallizes as an unconventional top-to-bottom dimer.

**Supplementary Fig. 10.**
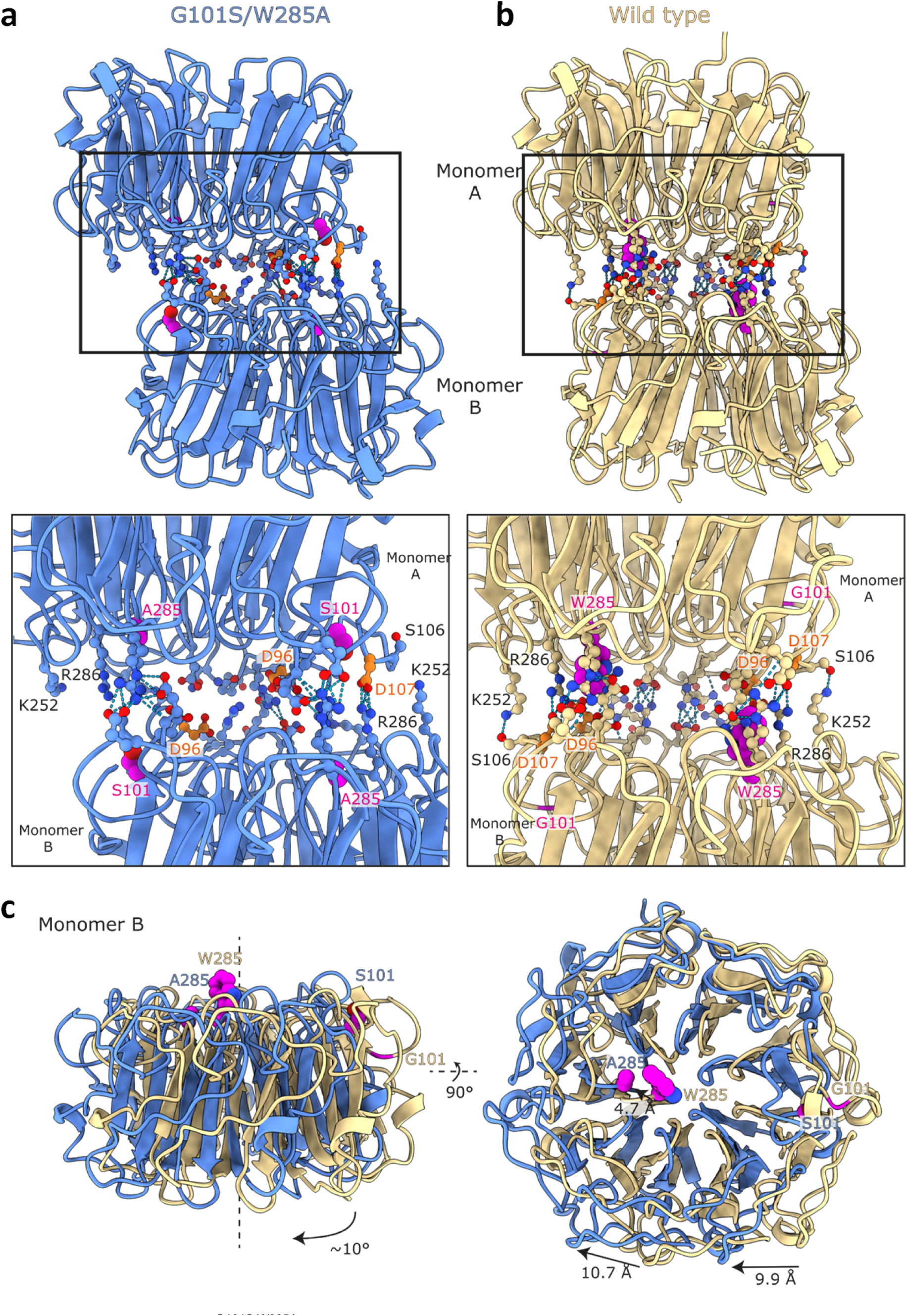
UVR8^G101S,W285A^ is an asymmetric dimer with an altered dimeric interface compared to wild-type UVR8. **a**,**b** A comparison of the UVR8^G101S,W285A^ (G101S/W285A) and UVR8 (wild type) homodimer orientation based upon the superposition of monomer A. Monomer B of UVR8^G101S,W285A^ is rotated relative to the wild type (see **c**). Both structures are depicted as ribbons (UVR8^G101S,W285A^, blue; wild-type UVR8, yellow). All side chains are depicted as ball-and-stick models. The sites of mutation residues 101 and 285 are colored in magenta. D96 and D107 are highlighted in orange. Hydrogen bonds or salt-bridges are colored in teal. The salt-bridges found in the symmetrical UVR8 dimer are no longer present in UVR8^G101S,W285A^. A list of interaction residues is shown in Supplemental Table 2. **c** Comparison of monomer B of UVR8^G101S,W285A^ and UVR8 homodimers based on the superposition of monomer A (as in **a**). Monomer B of UVR8^G101S,W285A^ is rotated ∼10° and shows shifts of up to 10.7 Å relative to monomer B of a UVR8 dimer.

**Supplementary Fig. 11.**
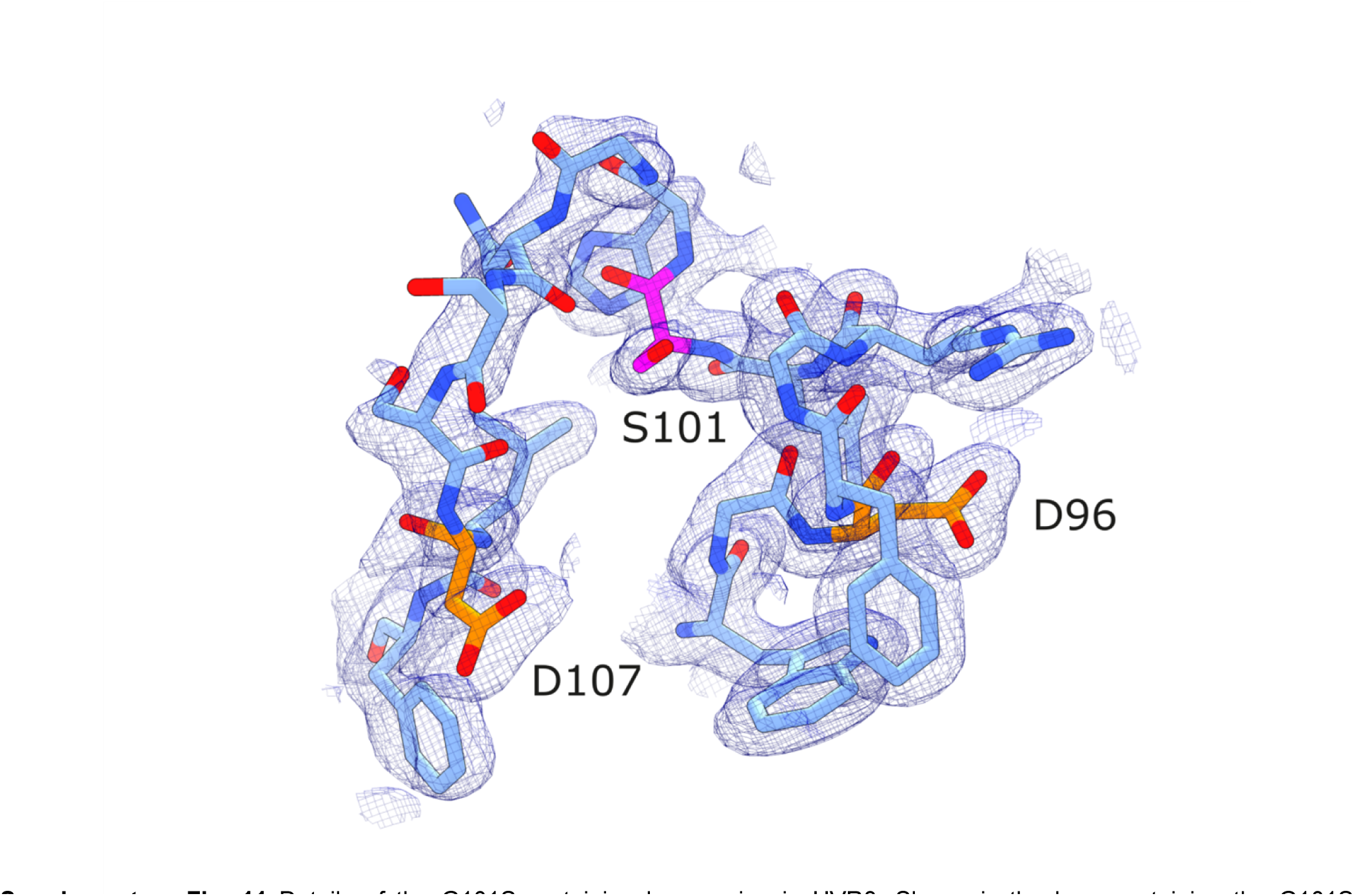
Details of the G101S-containing loop region in UVR8. Shown is the loop containing the G101S mutation from chain A of the UVR8^G101S,W285A^ structure (in bonds representation, in blue). A 2mFo-DFc electron density map contoured around all atoms depicted at a level of 1 σ is shown alongside (blue mesh). The site of mutation, G101S, is highlighted in magenta. Important residues D96 and D107 are highlighted in orange.

**Supplementary Fig. 12.**
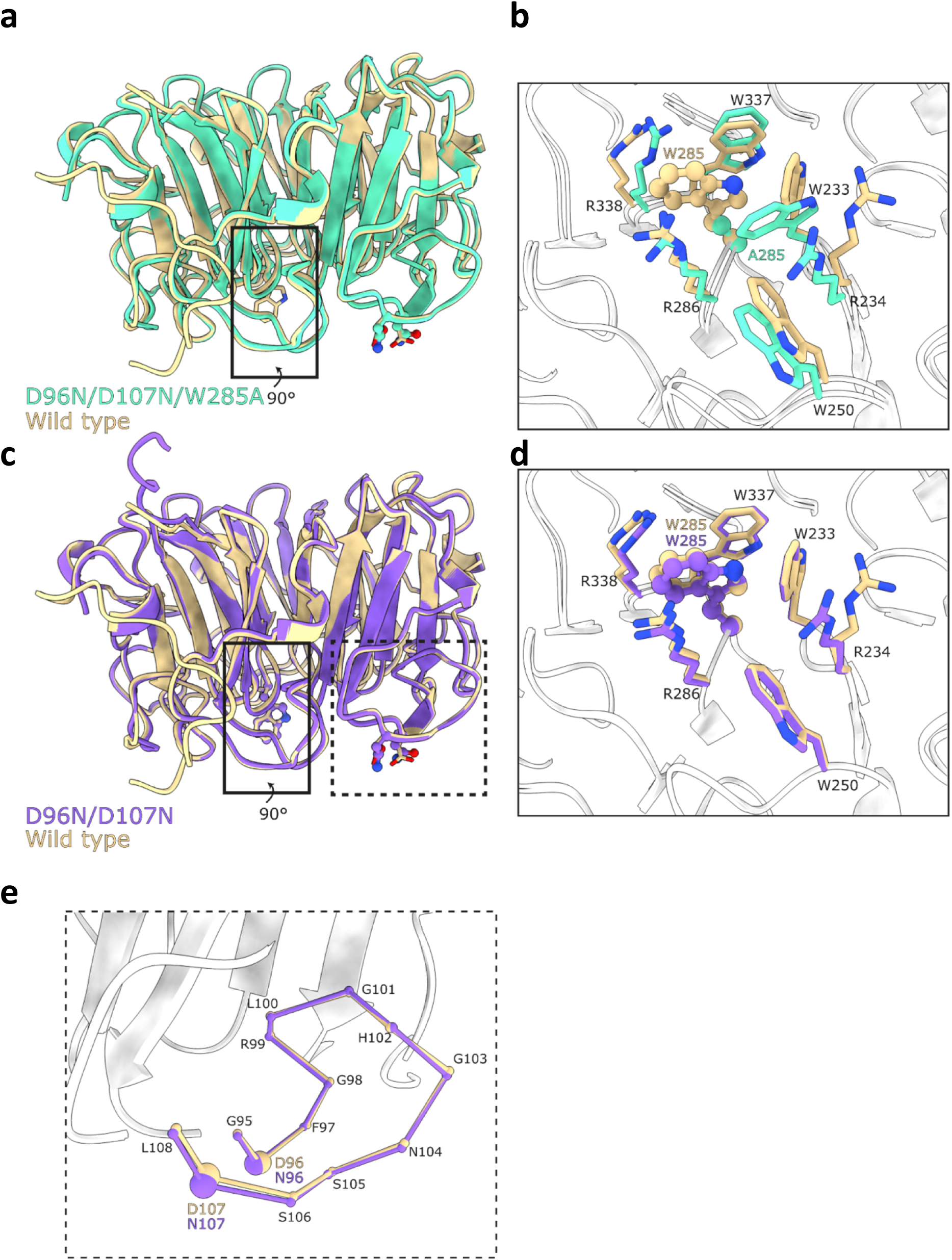
UVR8^D96N,D107N^ shows no major structural changes. **a**,**c** Superposition of UVR8^D96N,D107N,W285A^ (D96N/D107N/W285A; green) or UVR8^D96N,D107N^ (D96N/D107N; purple) with a wild-type UVR8 (wild type; yellow) in ribbon representation. The sites of mutation, residues 96, 107 and 285, are represented by a ball-and-stick model. **b**,**d** A zoomed in view of the site containing the W285A mutation. The site of mutation is represented as a ball-and-stick model and the surrounding residues are shown as sticks. **e** Zoomed-in view of the loop containing the D96N,D107N mutations. The loop is represented as a ball-and-stick model to highlight the loop with each ball corresponding to a Cα carbon.

**Supplementary Fig. 13.**
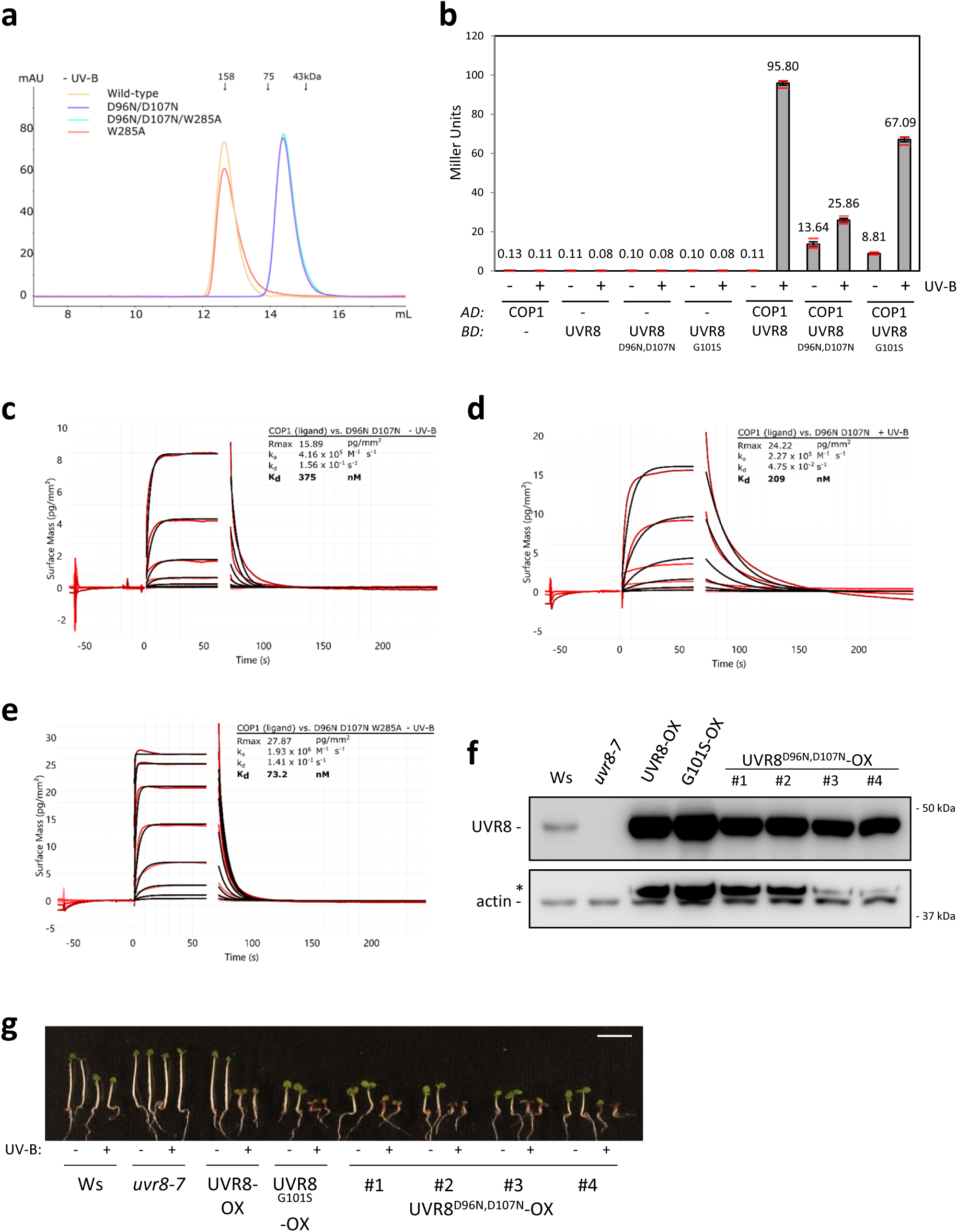
Lines overexpressing UVR8^D96N,D107N^ phenocopy UVR8^G101S^-overexpression lines. **a** Size-exclusion chromatography assay of recombinant UVR8 (wild-type), UVR8^D96N,D107N^, UVR8^W285A^, and UVR8^D96N,D107N,W285A^ proteins expressed in Sf9 insect cells. **b** Quantitative Y2H analysis of the interaction of COP1 with UVR8, UVR8^D96N,D107N^, and UVR8^G101S^ in the absence or presence of UV-B. AD, activation domain; BD, DNA binding domain. **c**–**e** Binding kinetics of the full-length UVR8^D96N,D107N^ and UVR8^D96N,D107N,W285A^ versus the COP1 WD40 domain obtained by GCI experiments. Sensorgrams of protein injected are shown in red, with their respective heterogenous ligand binding model fits in black. The following amounts were typically used: ligand, COP1 (2000 pg/mm^2^); analyte, UVR8 (2 μM highest concentration). ka = association rate constant, kd = dissociation rate constant, Kd = dissociation constant. **f** Immunoblot analysis of UVR8 and actin (loading control) protein levels in wild type (Ws), *uvr8-7*, *uvr8-7*/Pro_35S_:UVR8 (UVR8-OX), *uvr8-7*/Pro_35S_:UVR8^G101S^ #2 (G101S-OX), and four independent lines of *uvr8-7*/Pro_35S_:UVR8^D96N,D107N^ (UVR8^D96N,D107N^-OX #1–4). **g** Representative images of seedlings described in **f** grown for 4 d in white light or white light supplemented with UV-B. Bar = 5 mm.

**Supplementary Fig. 14.**
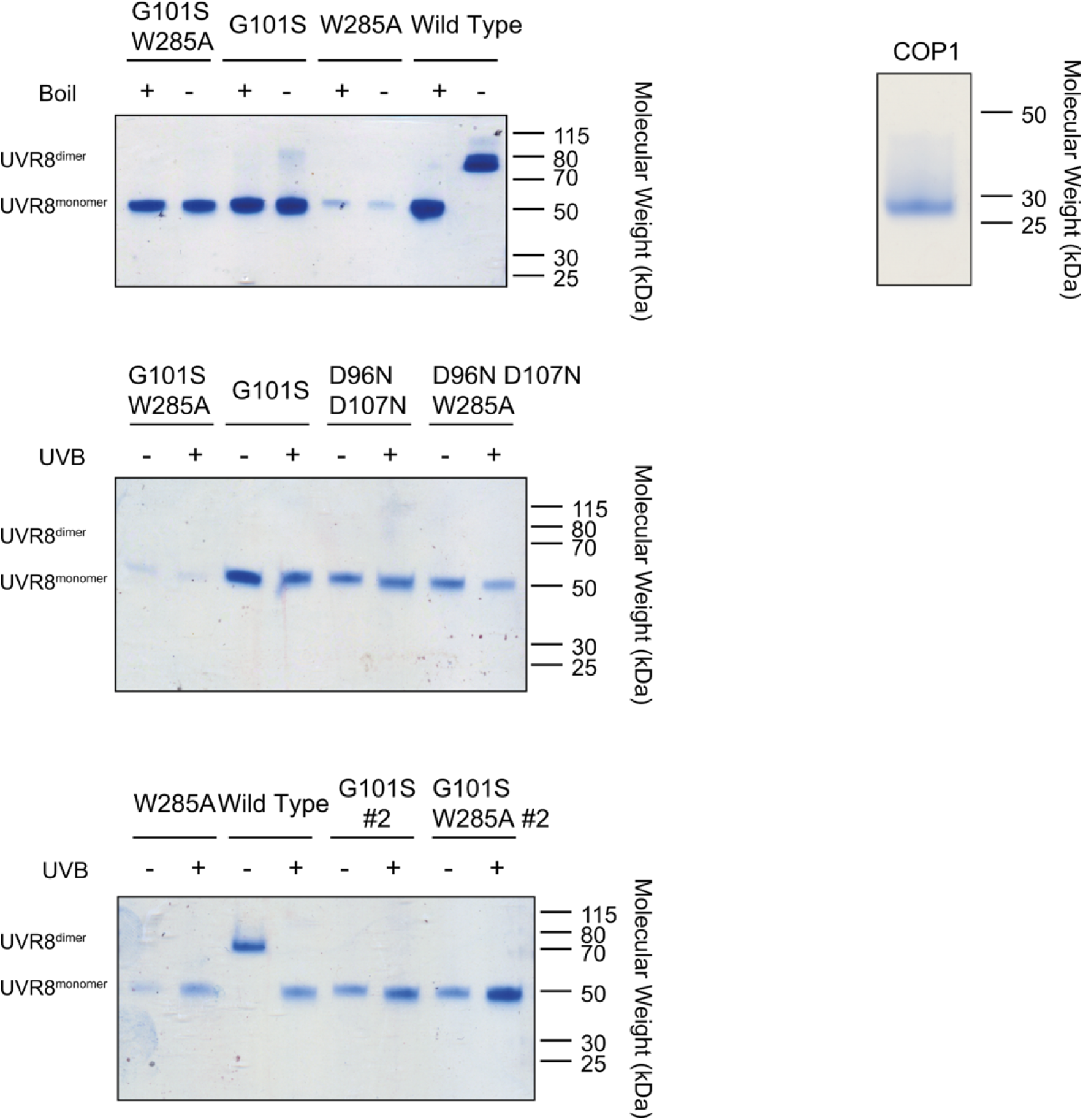
Coomassie-stained 10% SDS-PAGE gels of purified proteins show high purity. Representative SDS-PAGE gels of proteins used in this study. “#2” in the bottom panel indicates proteins from a second batch of purification.

**Table 1.**
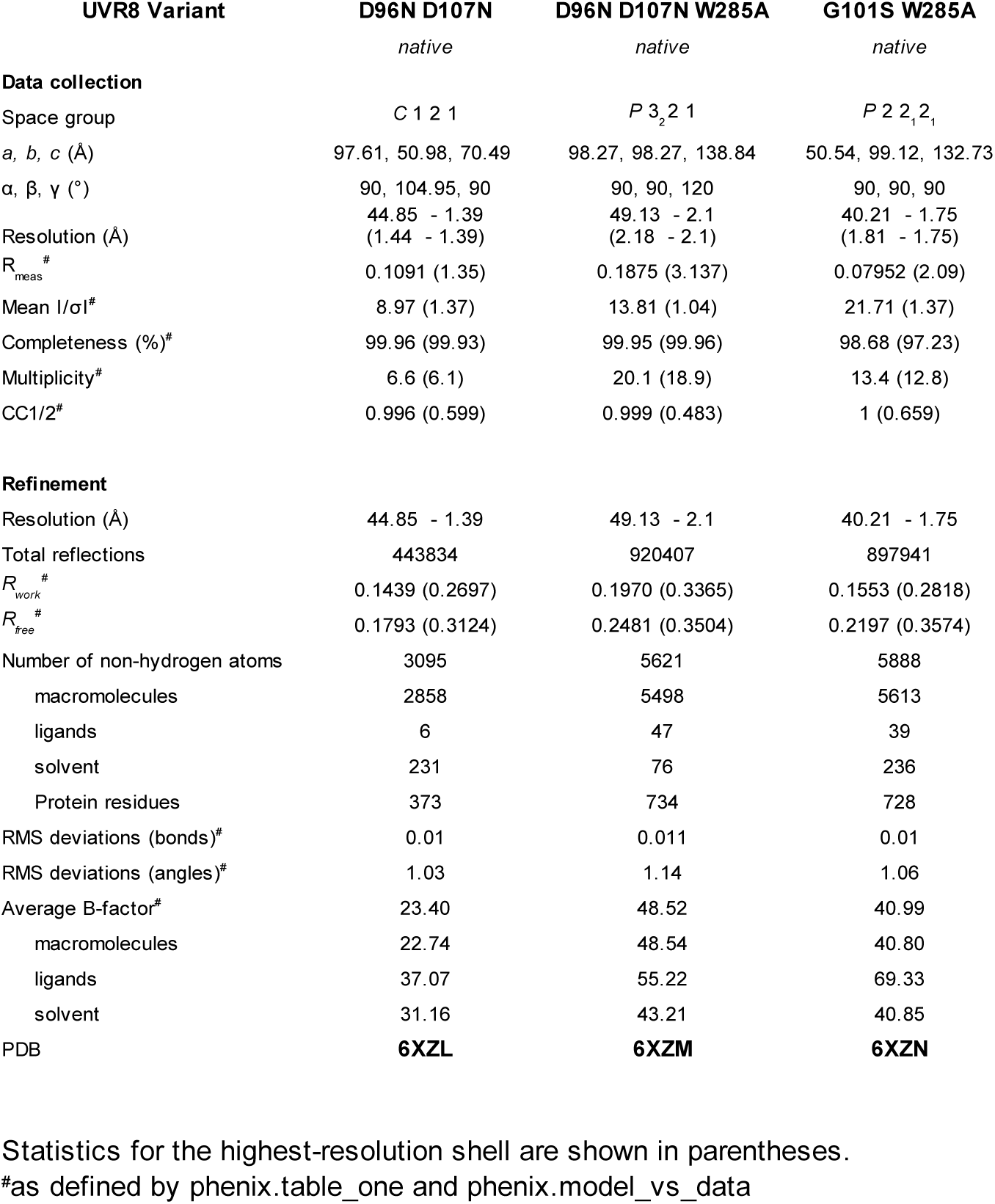
Data collection and refinement statistics.

**Supplementary Table 2:**
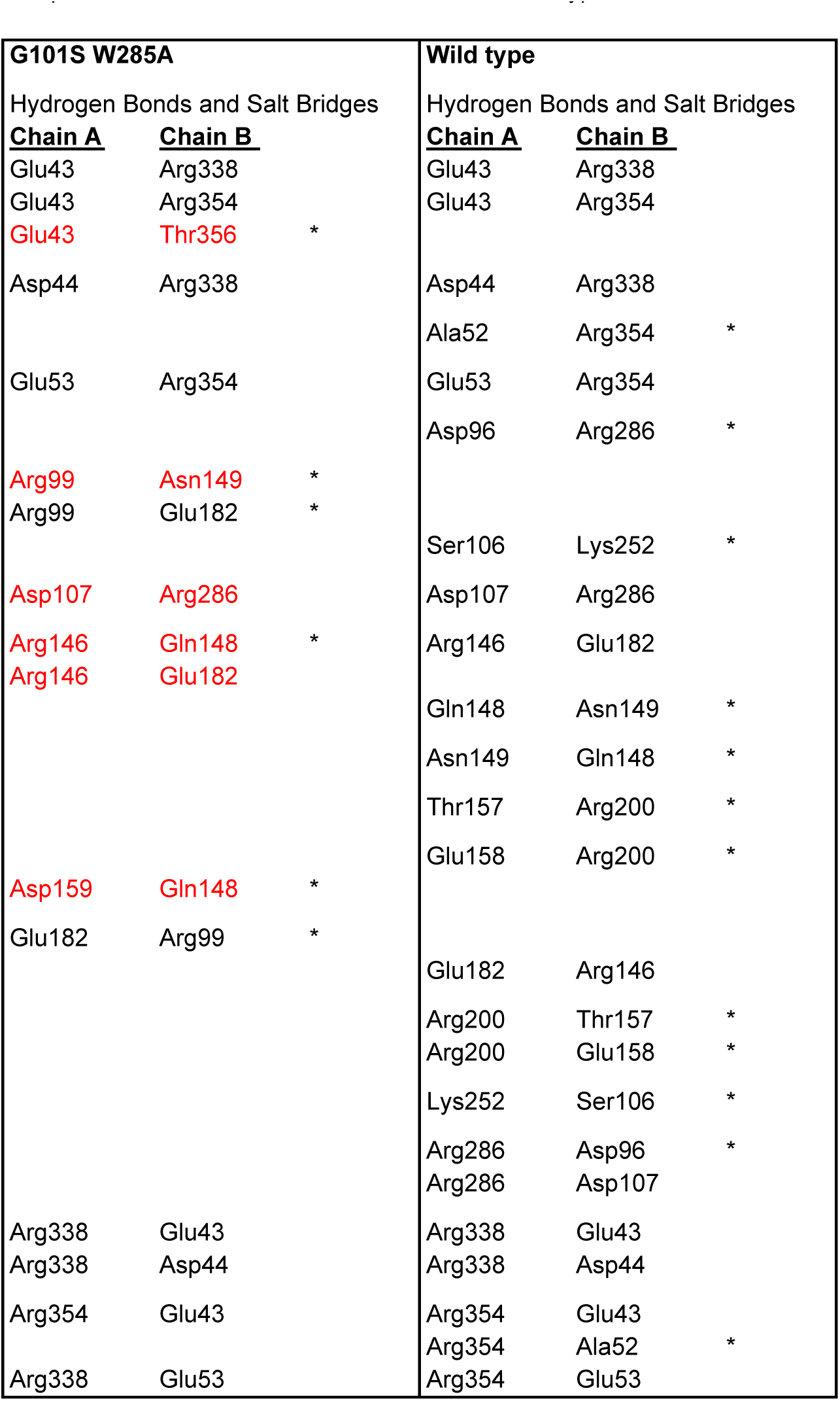
List of residues forming hydrogen bonds and salt-bridges at the UVR8 ^G101S,W285A^ and UVR8 dimer interface. Pairs colored black represent reciprocal interactions found between monomers of either UVR8 ^G101S,W285A^ or UVR8. Red pairs are non-reciprocal pairs between monomers. The wild-type UVR8 forms a symmetric dimer. Interaction pairs noted with an asterisk (*) denote interactions that are unique to the UVR8 ^G101S,W285A^ dimer or to the wild-type dimer.

